# Adenosine diphosphate contributes to wound healing in diabetic mice through P2Y_1_ and P2Y_12_ receptors activation

**DOI:** 10.1101/2020.10.22.350785

**Authors:** Paula A. Borges, Ingrid Waclawiak, Janaína L. Georgii, Janaína F. Barros, Vanderlei S. Fraga-Junior, Felipe S. Lemos, Thaís Russo-Abrahão, Elvira M. Saraiva, Christina M. Takiya, Robson Coutinho-Silva, Carmen Penido, Claudia Mermelstein, José R. Meyer-Fernandes, Fábio B. Canto, Josiane S. Neves, Paulo A. Melo, Claudio Canetti, Claudia F. Benjamim

## Abstract

Several studies have shown the importance of purinergic signaling in various inflammatory diseases. In diabetes mellitus, there is an increase in the activity of some nucleotidases suggesting that this signaling may be affected in the diabetic skin. Thus, the aim of our study was to investigate the effect of ADP on wound healing in diabetic skin. Swis and C57BL/6 mice were pharmacologic induced to type 1 diabetes and submitted to a full-thickness excisional wound model to evaluate the effect of ADP as a topic treatment. Adenosine diphosphate accelerated cutaneous wound healing, improved the new tissue formation, and increased collagen deposit by positively modulating P2Y_1_ and P2Y_12_ and TGF-β production. In parallel, ADP reduced reactive oxygen species production and TNF-*α* levels, while increased IFNγ, IL-10 and IL-13 levels in the skin. Also, ADP induced the migration of neutrophils, eosinophils, mast cells, TCRγ4^+^, and TCRγ5^+^ cells while reduced Treg cells towards the lesion at day 7. In accordance, ADP increased the proliferation and migration of fibroblast, induced myofibroblast differentiation and keratinocyte proliferation in a P2Y_12_-dependent manner. We provide the first evidence of ADP acting as a potent mediator on skin wound resolution and a possible therapeutic approach for diabetic patients worldwide.

## Introduction

Wound healing is a complex, dynamic and multi-mediated process characterized by a highly regulated cascade of events requiring the interaction of many cell types, including inflammatory and immune cells. Normal healing process occurs over a range of overlapping events: inflammation, granulation tissue formation, and remodeling. Impaired wounds are often associated with pathologic inflammation due to a persistent, incomplete, or uncoordinated healing process [1, 2].

Patients suffer from abnormalities of wound healing worldwide; in particular, in senescence, diabetes, ischemia, peripheral vascular disease, or cancer [3, 4]. Chronic wounds are reported to affect around 6.5 million patients just in USA; the estimated annually cost is more than US$ 25 billion for wound-related complications and healthcare system. In Brazil, the most populous country in Latin America, about 40 to 60% of non-traumatic lower limb amputations occur in diabetic patients; whereas, about 85% are related to foot ulcers [5-7].

ADP plays a pivotal role in the physiologic process of hemostasis and platelet aggregation. ADP activates P2Y_1_, P2Y_12_, and P2Y_13_ receptors, which are expressed by monocytes/macrophages, lymphocytes, mast cells, fibroblasts, keratinocytes, endothelial cells, eosinophils, platelets, neutrophils, and dendritic cells [8-10]. Neuroprotective function for ADP was demonstrated in zebrafish retina since it mitigated the excessive cell death and tissue damage; additionally, it stimulated cellular proliferation after injury [11]. This study aims to explore the role of ADP in accelerating wound healing in diabetic mice, considering that chronic wounds are a relevant health problem evidenced by the lack of an effective treatment, especially in diabetic patients, and also the pro-inflammatory, cell proliferative and pro-resolution effects of purinergic antagonists. The full-thickness excisional wound murine model was used to address our aim since it is the most reproducible and feasible model for tissue repair.

## Methods

### Mice

Male Swiss and C57BL/6 mice, obtained from the Institute of Science and Technology in Bio-models at Oswaldo Cruz Foundation, were used for full-thickness excisional wound models. For the cutaneous leishmaniasis lesion model, we used male BALB/c mice, obtained from the Microbiology and Parasitology Department animal facility at Biomedical Institute in Fluminense Federal University. All procedures described were approved by the Ethics Committee for the Use of Animals of the Federal University of Rio de Janeiro (CEUA/UFRJ: 093/15 and IMPPG 128/15).

### Induction of diabetes mellitus

Diabetes was induced by alloxan (65 mg/kg, i.v.) in mice fasted for 12 h [12,13]. Non-diabetic mice were injected with saline. Diabetes was confirmed 7 days later when blood glucose concentration was at least 350 mg/dl. The glucose levels were still elevated (over 350 mg/dl) at day 30 after alloxan injection.

### Full-thickness excisional wound model

At day 7 after alloxan or saline administration, mice were i.p. anesthetized (ketamine 112 mg/kg and xylazine 7.5 mg/kg) and a full-thickness excisional wound (10 mm in diameter) was executed on the dorsum using biopsy punch. Wounds were treated once a day for 5 or 14 days with topical application of 5’-AMP, ADP, ATP, adenosine, or pyrophosphate (Sigma-Aldrich, St Louis, MO) at 30 μM (30 μL - 15.4 µg/kg), or vehicle (30 μL of saline/mouse) or until the determined day for sample collection.

### Wound area quantification

To determine the wound closure rate, the wound area was evaluated at days 0, 3, 7, 10, and 14 after wounding. Photos were taken at a standard distance using a tripod and were analyzed using ImageJ software. Data were expressed as a percentage of the initial wound area.

### Treatments

The administration of Clopidogrel® (Clop - 5 mg/kg) was performed daily by oral gavage, 1 h before ADP administration, for 14 days. Treatments of P2Y_1_ (MRS 2179 - 30 µM/30 μL/mouse - Tocris, Bioscience, UK) and P2Y_12_ (MRS 2395 - 30 µM/30 μL/mouse - Sigma-Aldrich) receptor antagonists, and ATP diphosphohydrolase (apyrase - 6 U/mL, 30 μL/mouse - Sigma-Aldrich) were topically applied for 14 days, 30 min before ADP administration.

### Hematoxylin and eosin and total collagen quantification

Wounds tissues were paraffin-embedded and cut in 5-μm thick sections. Hematoxylin and eosin staining were performed, as previously described. Skin sample sections (7-μm) were stained with Picro-Sirius Red for total amount of collagen, as previously reported (Barros et al., 2019). The quantification was determined by morphometric analysis using a quantitative imaging software (ImagePro Plus, version 4.5.1). The percentage of collagen per field was obtained by dividing the total area by the fibrosis area.

### Ecto-nucleotidase activity

Ecto-nucleotidase activity was determined in the wound homogenates by the rate of inorganic phosphate (Pi) release using the malachite green reactions, as described elsewhere [14]. The concentration of Pi released in the reaction was determined by a Pi standard curve and expressed as nucleotidase activity (nmol Pi x h^-1^x mg ptn^-1^).

### Imunohistochemistry

Wound samples collected at day 7 were paraffin-embedded, sections (7-μm) cut and immunostained for several markers, as described previously [15]. The specific markers are detailed in the supplemental material. The data were expressed as number of positive cells per field. For collagen type markers, we developed a score method for the semi-quantification of collagen deposits performed by two different observers.

### Immunofluorescence

Wound sections (5-μm) were immunostained against *α*-SMA (A-2547, 1:200, Sigma-Aldrich) as previously reported [16]. Control experiments with no primary antibodies showed only faint background staining (data not shown).

### Cell proliferation

Primary neonate dermal fibroblasts (2 x 10^4^ cells) were used for BrdU staining proliferation assay, as previously described [17]. The images were captured using a fluorescence microscope and analyzed using ImageJ software. Results were expressed as the percentage of BrdU^+^ cells by total number of cells labeled with DAPI.

### Wound scratch assay

Primary dermal fibroblasts were seeded and grown until 90% confluence to evaluate migration-induced effect of ADP (10, 30, or 100 µM), as previously reported [18]. Pictures of the scratched areas were taken at 0, 6, 12, 18, and 24 h using an inverted microscope equipped with a digital camera (BEL Engineering - Monza, Italy). The areas were measured using the ImageJ software, and fibroblast migration expressed as % of area still open related to the initial area (0 h).

### Flow Cytometry

Flow cytometry of the wound tissues was performed as previously described [19]. Briefly, wound tissues were digested by an enzyme cocktail (supplemental material) and the cells (10^6^ cells/mL) were subjected to FACs procedure, stained, and analyzed. Lymphocyte populations recovered from skin and draining inguinal, axillary, and brachial lymph nodes were also analyzed. For skin Treg cell analysis, samples were enriched by Percoll gradient for mononuclear cells. Samples were acquired with BD FACS Canto II (BD Biosciences, San Jose, CA) and then analyzed with FlowJo software. Gating strategy and the list of antibodies are described in the supplemental material.

### Eosinophils and mast cells infiltrate

Skin sections (5-μm) were stained with modified Sirius Red or Alcian Blue for eosinophils and mast cells, respectively, as described elsewhere [20, 21]. Images were taken using a digital camera coupled to the microscope (Olympus BX53) at 40x magnification. Twenty fields were analyzed per wound/animal (n=3) and the data were expressed as number of eosinophils or mast cells/mm^2^.

### Myeloperoxidase activity

The number of neutrophils was indirectly determined by myeloperoxidase enzyme activity in the wounds removed 7 days after wounding, as previously described [22]. Neutrophils numbers were estimated by a standard curve, using neutrophils obtained 6 h after i.p. administration of 3% thioglycollate (>90% of neutrophils). Proteins were measured by the Bradford method. Results were expressed as number of neutrophils/mg of protein.

### ELISA

Cytokine quantification was performed in wounds obtained at day 7 using PeproTech kits following manufacturer’s instructions. The results were expressed as pg or ng of cytokine/mg of protein.

### Superoxide assay

The superoxide production assay was performed by the nitroblue tetrazolium (NBT) reaction with reactive oxygen species resulting in formazan as final product [23]. Briefly, the wounds were removed at day 3 and homogenized in PBS containing protease inhibitors for the assay. The formazan formed was measured by ELISA reader (620 nm, Spectra Max-250, Molecular Devices). Results were expressed as µg of formazan/mg of protein.

### Cytometric Bead Array (CBA)

Cytokine concentration in the wounds was determined by flow cytometry using the kit CBA Mouse Inflammation (BD Biosciences, San Diego, CA), following manufacturer’s instructions. This CBA kit allows measurements of IL-6, IL-10, C-C motif chemokine ligand 2, IFN-γ, TNF-*α*, and IL-12p70. Sample processing and data analysis was acquired by FACS Calibur flow cytometer (BD Bioscences) and FCAP Array software, respectively. Results were expressed as pg or ng of cytokine/mg of protein.

### Western blotting

Wound homogenates (30 mg of protein) collected at day 7 were prepared as previously described [24]. Immunoreactive bands for α-SMA (1:1000 - Sigma-Aldrich) and β-actin (1:1000 - Cell Signaling, Danvers, MA) were visualized using an enhanced chemiluminescence reagent (Amersham ECL, Biosciences) and pictures were recorded using Healthcare ImageQuant LAS 4000 (GE Healthcare Life Sciences). Densitometry analysis was performed using ImageJ software and the results expressed as the ratio of α-SMA/β-actin (housekeeping).

### Statistical analysis

Statistical differences in the wound closure experiments were determined using two-way ANOVA with the Bonferroni post-test using the GraphPad Prism software. The significance of other experiments was determined by unpaired Student’s t test.

## Results

### ADP improves wound healing in diabetic mice

Swiss male diabetic and non-diabetic mice were topically treated with saline or ADP 30 μM (30 μL - 15.4 µg/kg), every day for 5 days after wounding. ADP was effective in accelerating the wound closure in diabetic mice, but without changing the wound healing in non-diabetic mice (Figure 1a and 1b). ADP-treated diabetic mice presented 60 % wound closure versus 2 % in saline-treated mice at day 7. More importantly, the wound closure profile of the diabetic animals treated with ADP was similar to saline-treated non-diabetic mice. Moreover, the only effective dose able to accelerate wound closure was 30 μM. Indeed, higher ADP doses seemed to delay wound healing observed until day 14 (Figure 1c).

**Figure 1.**
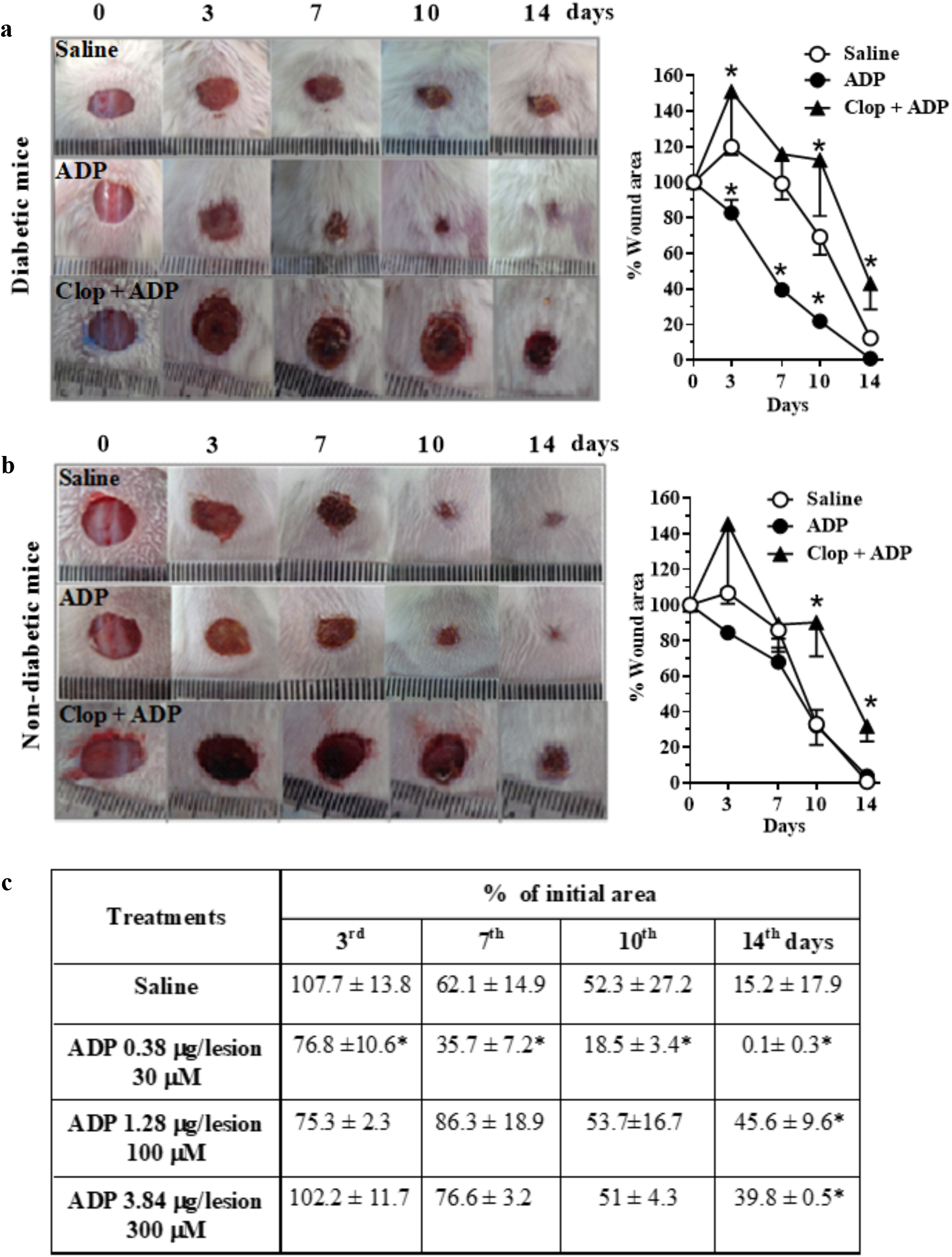
ADP accelerates wound healing in diabetic mice via P2Y_12_. Representative images and graphs of diabetic (**a**) and non-diabetic (**b**) mice that were submitted to excisional full-thickness wounding, and then, topically treated with ADP 30 μM (30 μL - 15.4 µg/kg) or saline every day for 5 days. One group of mice was treated by gavage with Clop (5 mg/kg) 1 h before ADP and saline, both once a day for 14 days. Open wound area was measured at days 0, 3, 7, 10 and 14. The areas at day 0 was considered 100 %, and the subsequent areas measured at different time-points were calculated as percentages (%) of the initial value. (**c**) Dose-effect curve of ADP treatment followed at days 3, 7, 10, and 14 after wounding. Data are expressed as mean ± standard error of the mean. *P<0.05 by Student’s t test, compared to saline-treated diabetic mice; n=7-10 per group. Panels A and B are representative of three or more experiments; panel C represents one experiment.

### Clopidogrel (Clop) impairs ADP-induced wound closure

To assess the role of P2Y_12_ and ADP in our model, a P2Y_12_ irreversible antagonist, Clop (5 mg/kg), impaired the ADP-treated wound closure in diabetic mice. It was characterized by an increase of the lesion size and a worsening of the wound general aspects, at all the time-points evaluated (Figure 1a). Furthermore, Clop administration also worsened the saline-treated wound of diabetic mice. Still, an endogenous critical role of ADP/P2Y_12_ for tissue repair was suggested since Clop treatment also impaired saline- or ADP-treated wounds of non-diabetic mice (Figure 1b and Supplemental Figure 1).

Assuming that ADP is the major antagonist of P2Y_12_ [25,26], our data demonstrates an unequivocal proof of ADP’s role in accelerating wound closure of diabetic mice.

### P2Y_1_ is also involved in ADP effects

ADP receptors involvement in wound healing was verified by another P2Y_12_ antagonist (MRS2395) and by a P2Y_1_ antagonist (MRS2179), both at 30 μM/mouse. Both antagonists impaired the wound closure induced by ADP until day 7; however, at day 10 and 14 the wound healing profile was identical to that of ADP-treated group (Figure 2a-b). P2Y_1_ and P2Y_12_ antagonists alone did not alter the wound closure in diabetic mice. We observed in the Figure 1 that Clop treatment not only impaired the beneficial ADP effect, but also worsened the healing process, and we expected to observe the same response with MSR2395. However, the administration routes were different between these drugs, and Clop is an irreversible antagonist, unlike the MRS2395 and MRS2179 that are competitive antagonists.

**Figure 2.**
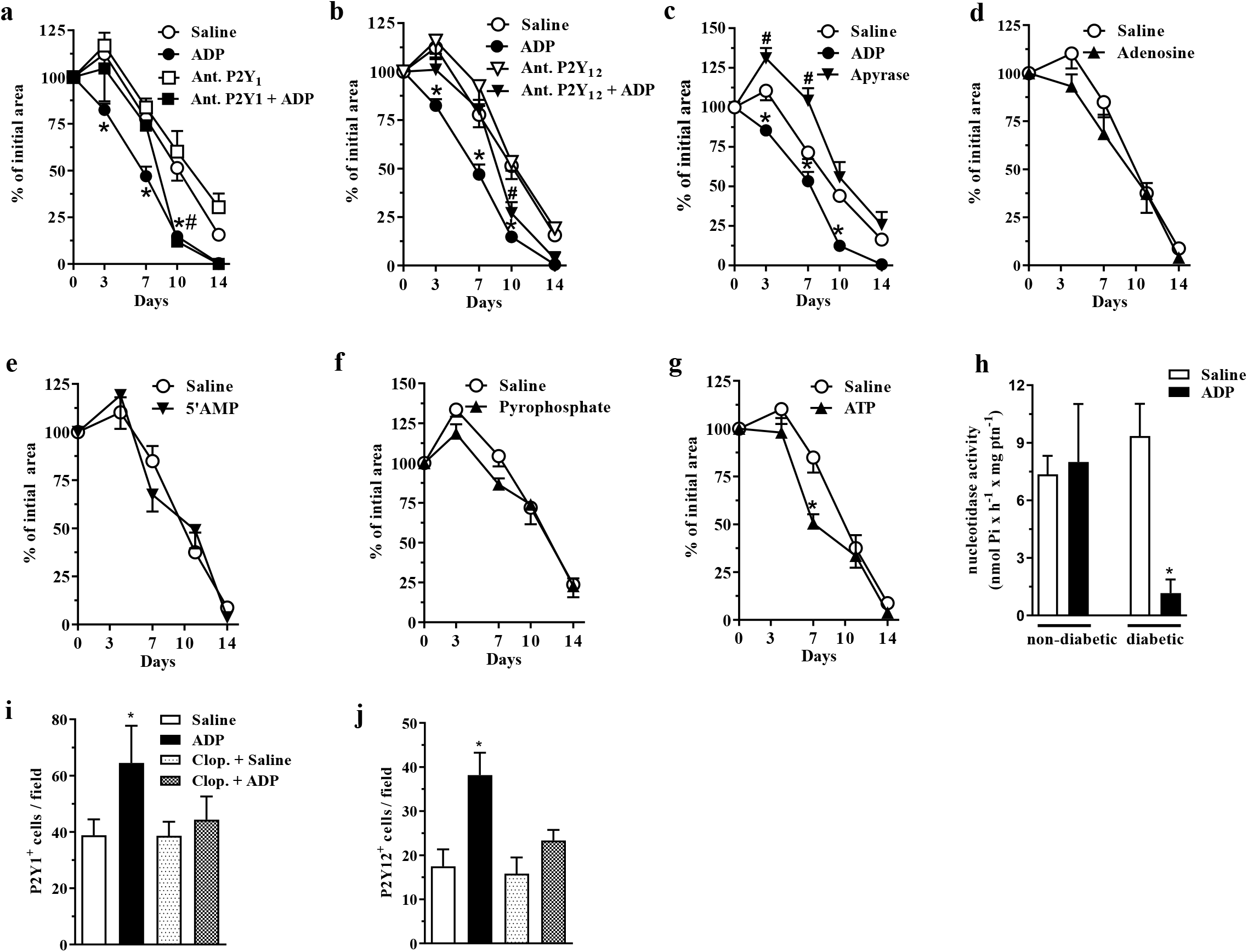
Central role of ADP, P2Y_1_ and P2Y_12_during wound healing of diabetic mice. (**a-b)** Diabetic mice were submitted to excisional full-thickness wounding and, then, topically treated with P2Y_1_ or P2Y_12_ antagonists (30 µM/mouse - 30 µL) 30 min before ADP (30 μM/mouse) or saline administration. Both antagonists were applied every day for 14 days. Open wound area was followed over time as described in Figure 1. ^*^P<0.05 by Student’s t test, compared between saline-or ADP-treated wound, ^**#**^P<0.05 by Student’s t test compared between P2Y_1_ or P2Y_12_ antagonists and saline-treated group, n=7-10 per group. (**c-g**) Diabetic mice were submitted to excisional full-thickness wounding and then topically treated with apyrase (6 U/mL), ATP, ADP, 5’AMP, adenosine, pyrophosphate (30 µM/mouse - 30 µL) or saline every day for 14 days. Open wound areas were followed over time as described in Figure 1. Data are expressed as mean ± standard error of the mean. ^*^P<0.05 by Student’s t test, compared to saline-treated group; ^**#**^P<0.05 by Student’s t test compared between apyrase- and saline-treated groups, n=8-10 per group. (**h**) Non-diabetic and diabetic mice were submitted to excisional full-thickness wounding and then topically treated with ADP (30 µM/mouse) or saline every day for 7 days. The nucleotidase activity was evaluated in the wound harvested at day 7. ^*^P<0.05 by Student’s t test, compared to saline-treated diabetic group, n=6 per group. (**i-j**) Diabetic mice submitted to excisional full-thickness wounding were topically treated with ADP (30 µM/mouse) or saline every day for 7 days. Both groups were treated by gavage with Clop (5 mg/kg) 1 h before saline or ADP treatment of the wounds. The wounds tissues were harvested at day 7 and stained for P2Y_1_ and PY_12_. The number of P2Y_1_^+^ and PY_12+_ cells per field was represented by the graphs. Data are expressed as mean ± standard error of the mean. ^*^P<0.05 by Student’s t test, compared to saline-treated group, n=6 per group.

### Apyrase worsens wound healing

Apyrase removes the γ-phosphate from ATP and the β-phosphate from ADP, yielding AMP [27]. Apyrase treatment worsened wound healing of diabetic mice, compared to saline- or ADP-treated diabetic wound (Figure 2c), confirming the crucial role of this nucleotide in tissue repair.

### Different nucleotides do not accelerate the wound healing

In order to certificate that ADP is the nucleotide responsible for the observed effect in our model, adenosine, 5’AMP, pyrophosphate or ATP were topically applied on the wounds of diabetic mice at 30 μM/mouse, the same ADP concentration used. Among the nucleotides tested only ATP showed a slight improvement of wound closure at day 7 (Figure 2d-g).

### ADP reduces ecto-nucleotidase activity in the wounds of diabetic mice

We tested a possible enzyme deregulation related to ADP degradation during diabetes. The ecto-nucleotidase activity observed in the ADP-treated wounds obtained from diabetic mice was reduced compared to saline-treated wounds from diabetic mice, and also when compared to both non-diabetic mice groups (Figure 2h). The same profile was observed in blood samples obtained from ADP-treated diabetic mice (data not shown). It seems that ADP treatment down-regulates ecto-nucleotidase activity only in diabetic mice, which seems to favor wound healing.

### ADP increases P2Y_1_^+^ and P2Y_12_ ^+^cells in

ADP-treated diabetic wounds presented higher expression of P2Y_1_ and P2Y_12_ at day 7, compared to saline-treated wounds. Clop treatment impaired ADP-induced P2Y_1_ and P2Y_12_ expression, whereas it did not change the receptors expression in saline-treated diabetic mice (Figure 2i-j).

### ADP improves tissue formation

Saline-treated wounds of diabetic mice presented edematous dermis, leukocyte infiltration (predominantly by mononuclear cells), and null (or partial) formation of epidermis at day 7. In the reticular dermis, exuberant granulation tissue formation and congested neovessels were observed (Figure 3a). Interestingly, ADP-treated wounds presented a chronological change of the regenerative process. The epidermis was regenerated and integrated to the underlying dermis, with hyperplasic suprabasal layers, and hyperkeratosis. In the dermis, there was an exuberant granulation tissue with inflammatory cell infiltrate comprising eosinophils, mast cells, myeloid progenitors, neutrophils, and mononuclear cells. Clop administration impaired tissue regeneration in saline-treated wounds, where denuded epidermis areas, necrotic dermis with an inflammatory infiltrate composed predominantly of polymorphonuclear cells, striking bleeding, and the absence of granulation tissue were observed. Clop administration also impaired wound healing in ADP-treated mice; however, with milder effects. In this case, wounds displayed a more organized reticular dermis, with collagen bundles parallel to the skin surface and interspersed with fibroblasts; a few vessels and inflammatory infiltrate were also noticed (Figure 3a).

**Figure 3.**
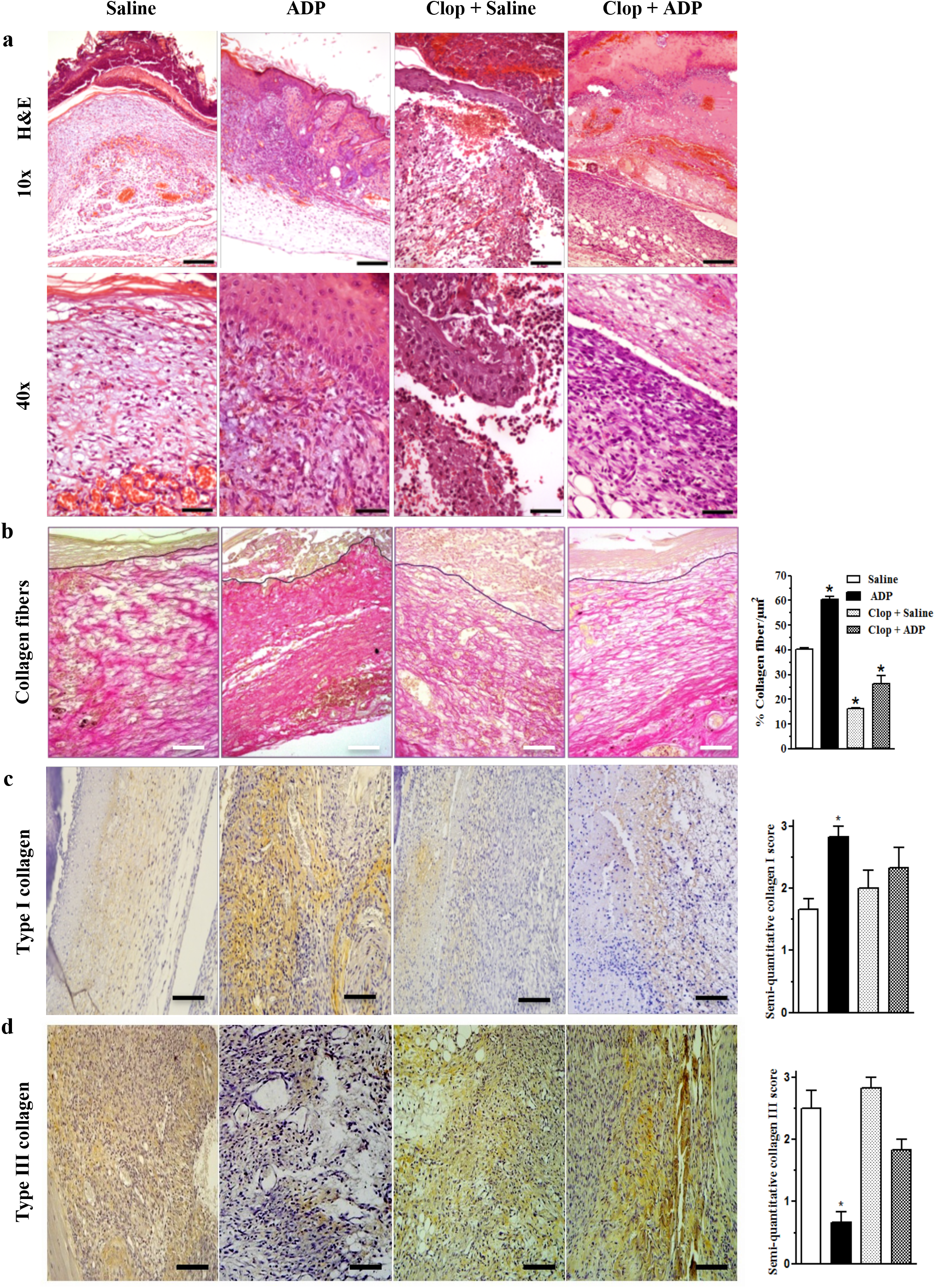
APD-treated wounds present an improved tissue repair and an increased collagen deposition. (**a**) Diabetic mice submitted to excisional full-thickness wounding were treated by gavage with Clop (5 mg/kg) 1 h before ADP (30 µM/mouse) or saline treatment, once a day for 7 days. Wounds were harvested at day 7 and stained with hematoxylin and eosin. Images representative of 4-5 mice per group. Scale bars: 10x=200 µm; 20x=100 µm; 40x=50 µm. (**b**) Collagen deposit (red staining) in wounds at day 7 stained with Picro-Sirius Red and the representative images are shown; bar graph summarized data from 5-6 mice per group, representative of three independent experiments. Scale bars: 50 µm. *P<0.05 by Student’s t test, compared to saline-treated group. (**c-d**) Type I and type III collagen staining by IHC. Scale bar 50 µm. Graphs with semi-quantification score for type I and type III collagen deposit. Data referred of one experiment with 3-4 mice per group.

Picro Sirius Red-stained photomicrographs (red staining) showed higher collagen fibers deposition in ADP-treated wounds of diabetic animals compared to the saline-treated wounds. Clop administration impaired collagen deposit in both ADP and saline-treated wounds. Collagen fibers quantification confirmed that ADP treatment enhanced collagen deposition while Clop administration impaired its accumulation (Figure 3b). ADP seemed to accelerate the switch of type III to type I collagen, a more mature fiber. Nevertheless, Clop administration reduced type I collagen deposit, without changing type III collagen production in both saline- and ADP-treated wounds (Fig. 3c-d). The results depicted in the bar graphs represent the photomicrographs.

### ADP induces keratinocyte proliferation

We also evaluated if ADP enhances re-epithelization. At day 7 after wounding, ADP-treated wounds presented higher number of Ki67^+^ cells, a cell proliferation marker, in the layer adjacent to the basal membrane when compared to saline-treated wounds. At day 14, the Ki67^+^ cells in ADP-treated wounds were reduced but still higher than in saline-treated wounds. Corroborating this result, epidermis area is also larger in ADP-treated wounds at day 7 compared to saline-treated wounds, while at day 14 it returned to normal (Figure 4a). The percentage of proliferating cells and the area of the epidermis, observed at day 7 and 14 post wounding, are shown in bar graph (Figure 4a).

**Figure 4.**
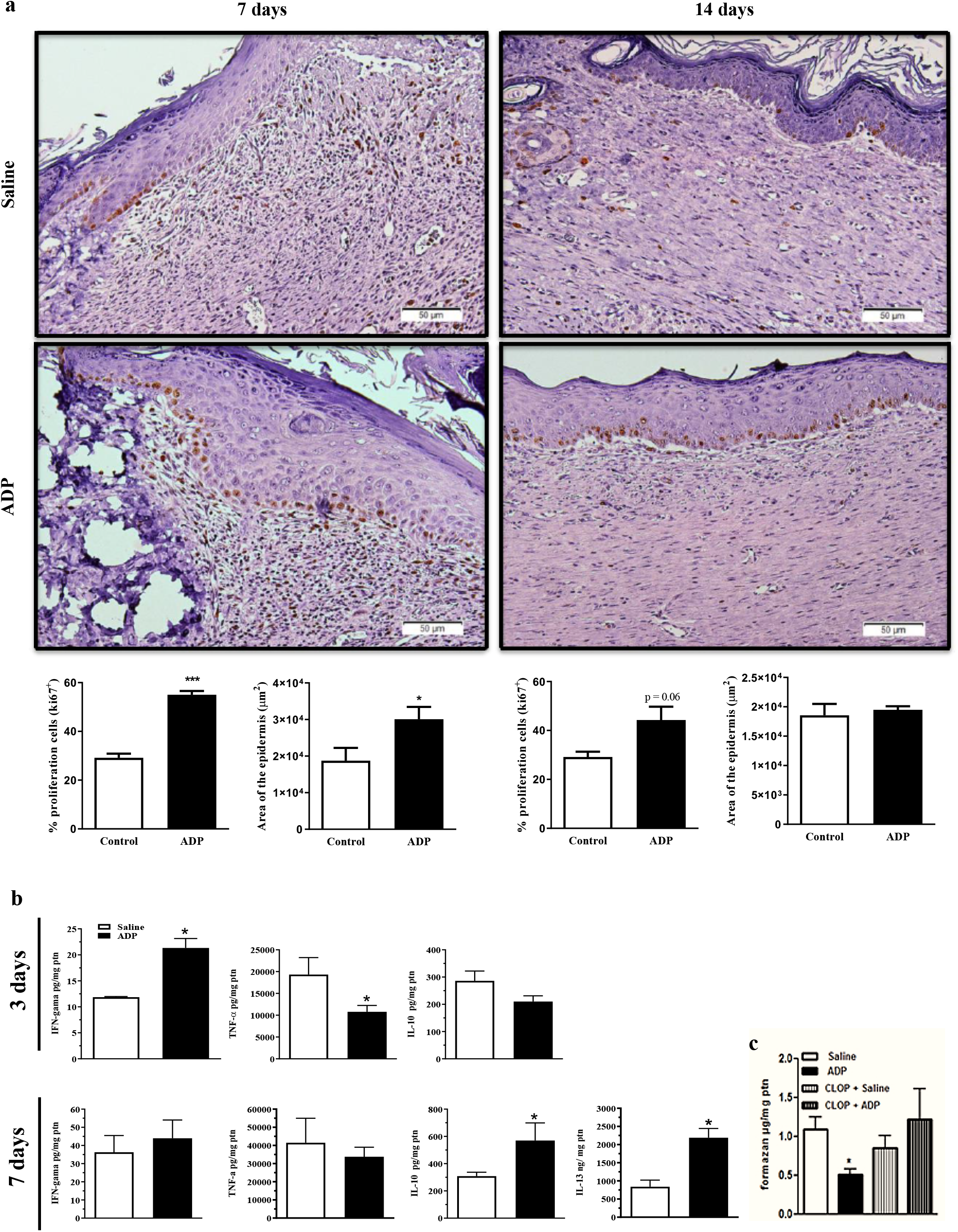
APD induces keratinocyte proliferation and modulates cytokine and free-radical production. Diabetic mice were submitted to excisional full-thickness wounding, and, then, topically treated with ADP (30 µM/mouse) or saline every day for up to 14 days. (**a**) Wounds were harvested from diabetic mice at days 7 and 14 after wounding, and stained for Ki67 by IHC, hematoxylin and eosin. Scale bars: 200 µm and 50 µm. The percentage of proliferating keratinocytes (Ki67^+^) and the area of epidermis were represented in bar graphs. Data are expressed as mean ± standard error of the mean. ^***^P<0.001 by Student’s t test, compared to saline-treated group; n= 6 per group. (**b**) Wounds were harvested from diabetic mice at days 3 and 7 after wounding and cytokine levels were evaluated by ELISA at day 3 and 7 after wounding. *P<0.05 by Student’s t test, compared to saline-treated group, n=4-6 per group. (**c**) Superoxide radical production was indirectly evaluated in the wounds obtained at day 7 after wounding by formazan generation as final product. Some animals were treated by gavage with Clop (5 mg/kg) 1 h before ADP or saline wound topic treatment. ^*^P<0.05 by Student’s t test, compared to saline-treated group, n=4-5 per group.

### ADP modulates the inflammatory response

At day 3 after wounding, ADP promoted increased INF-γ and reduced TNF-α levels without affecting IL-10 levels, while increased IL-10 and IL-13 levels were observed at day 7 (Figure 4b). No differences were detected in IL-6, IL-12p70 and C-C motif chemokine ligand 2 (CCL2/MCP-1) levels between groups (data not shown). We also observed reduction of reactive oxygen species production at day 7 post wounding after ADP treatment, while Clop administration restored reactive oxygen species production (Figure 4c), suggesting again the participation of P2Y_12_. These results suggest that ADP treatment controls inflammatory response associated with pro-resolution effects.

### ADP increases myofibroblasts population and TGF-β production

Myofibroblasts present high ability of promoting extracellular matrix protein production and wound contraction [1]. We observed that ADP treatment increased α-smooth muscle actin (α-SMA) expression in the dermis, which was reduced by Clop, while in Clop+ADP group a less dramatic reduction of myofibroblasts was observed (Figure 5a). ADP effect on α-SMA increased expression was also confirmed by WB assay (Figure 5b). In accordance, ADP at 30 µM also induced fibroblasts proliferation (Figure 5c) and migration (Figure 5d-e) in *in vitro* assays, suggesting a dose-dependent effect of ADP in wound healing.

**Figure 5.**
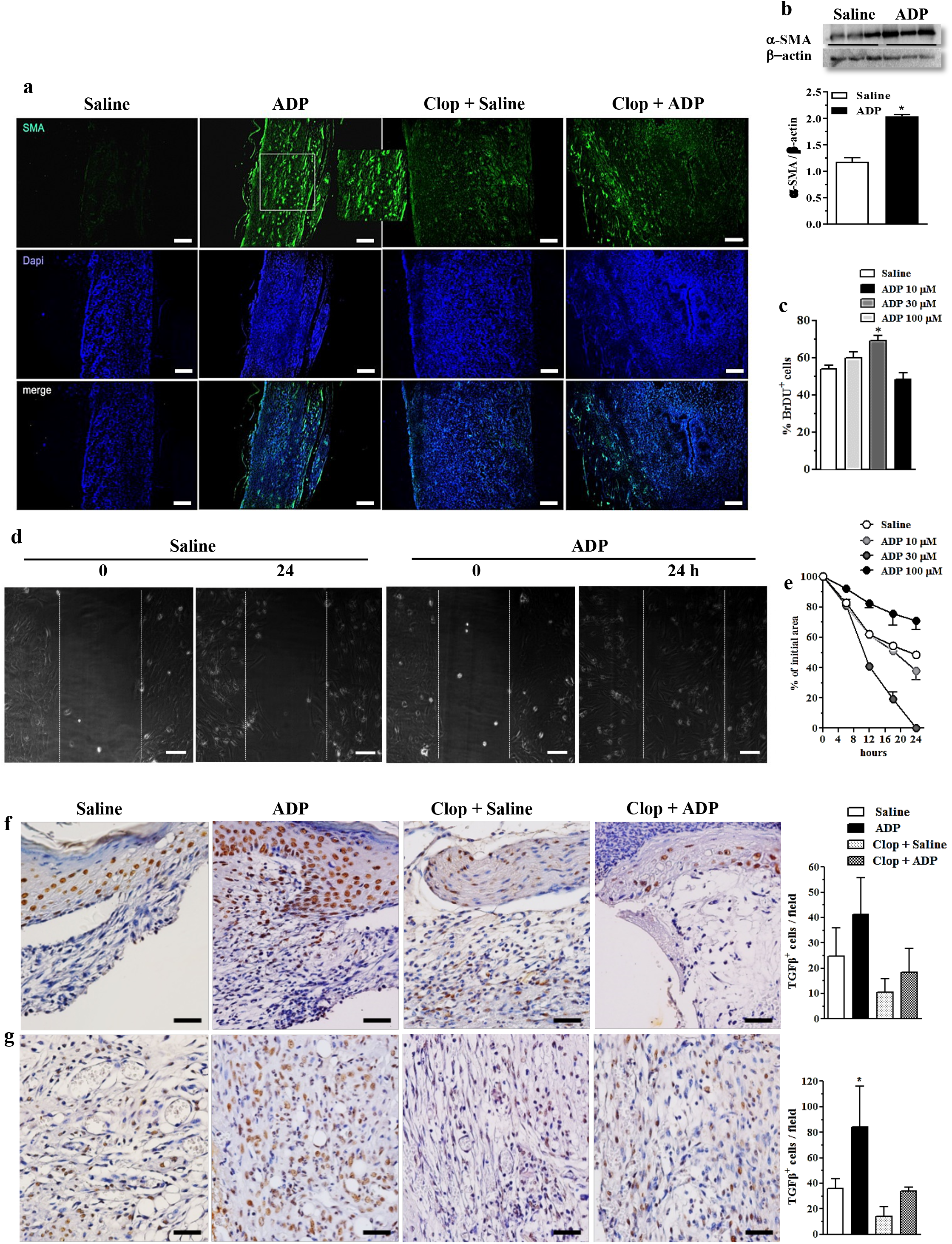
ADP activates myofibroblasts/fibroblasts and increases the amount of TGF-β^+^cells in the wounds of diabetic mice. (**a**) Diabetic mice were submitted to excisional full-thickness wounding and, then, topically treated with ADP (30 µM/mouse) or saline every day for 7 days. Some mice were treated by gavage with Clop (5 mg/kg) 1 h before ADP or saline administration, once a day for 7 days. Wounds harvested at day 7 were stained for *α*-SMA (green) and DAPI (blue) and analyzed by immunofluorescence. (**b**) Gel bands and graphs depicting the semi-quantification of *α*-SMA by WB at the same conditions in panel A. Each bar represents a pool of skin-derived protein extracts obtained from at least 5 mice. ^*^P<0.05 by Student’s t test compared to saline-treated group; data are representative of two independent experiments. (**c**) Primary culture of neonate murine dermal fibroblasts was plated for 24 h, incubated with BrdU for more 24 h and the cell proliferation was evaluated by immunofluorescence. ^*^P<0.05 by Student’s t test, compared to saline-treated group; data are representative of three independent experiments. (**d-e**) Primary dermal murine fibroblasts were plated for 24 h, pre-incubated with mitomycin-C 5 μg/mL for 2 h and then incubated with different concentrations of ADP. The open area between the front edges of the scratch were evaluated at 0, 6, 12, 18 and 24 h after scratch and expressed as % of initial area. Fibroblast culture images (D) represent only the first and last time point evaluated for cell migration. ^*^P<0.05 by Student’s t test, compared to saline-treated group, the data are representative of three independent experiments. (**f**) Photomicrographs and bar graph of TGF-β^+^ cells determined by immunohistochemistry in the epidermis and (**g**) dermis obtained at day 7 after wounding. ^*^P<0.05 by Student’s t test, compared to saline-treated group, n=8 per group.

TGF-β is a pivotal cytokine in wound healing [1], and ADP treatment seemed to increase TGF-β production by keratinocytes (Figure 5f – epidermis) and to increase the number of TGF-β^+^ cells in the dermis (Figure 5g).

### ADP treated wounds present a different leukocyte profile

Unbalance of numbers and/or activation of local leukocytes are common under diabetes condition, which compromises tissue repair [28]. Interestingly, we observed an increase of neutrophil recruitment in ADP-treated wounds (Figure 6a upper line and 6b). In parallel, a decrease in the inducible nitric oxide synthase^+^ cells and an increase in the arginase^+^ cells were detected in ADP-treated wounds (Figure c-d). This suggests that macrophage population switched towards an alternative-activated phenotype instead of an alteration in frequency, since F4/80^+^ macrophage numbers were similar between groups (Figure 6a bottom line). Clop treatment did not interfere with the change of macrophage phenotype in the wound induced by ADP (Figure 6c-d). Our data suggests an ADP-mediated, P2Y_12_-independent, skewed response towards pro-resolution scenario in the context of tissue injury.

**Figure 6.**
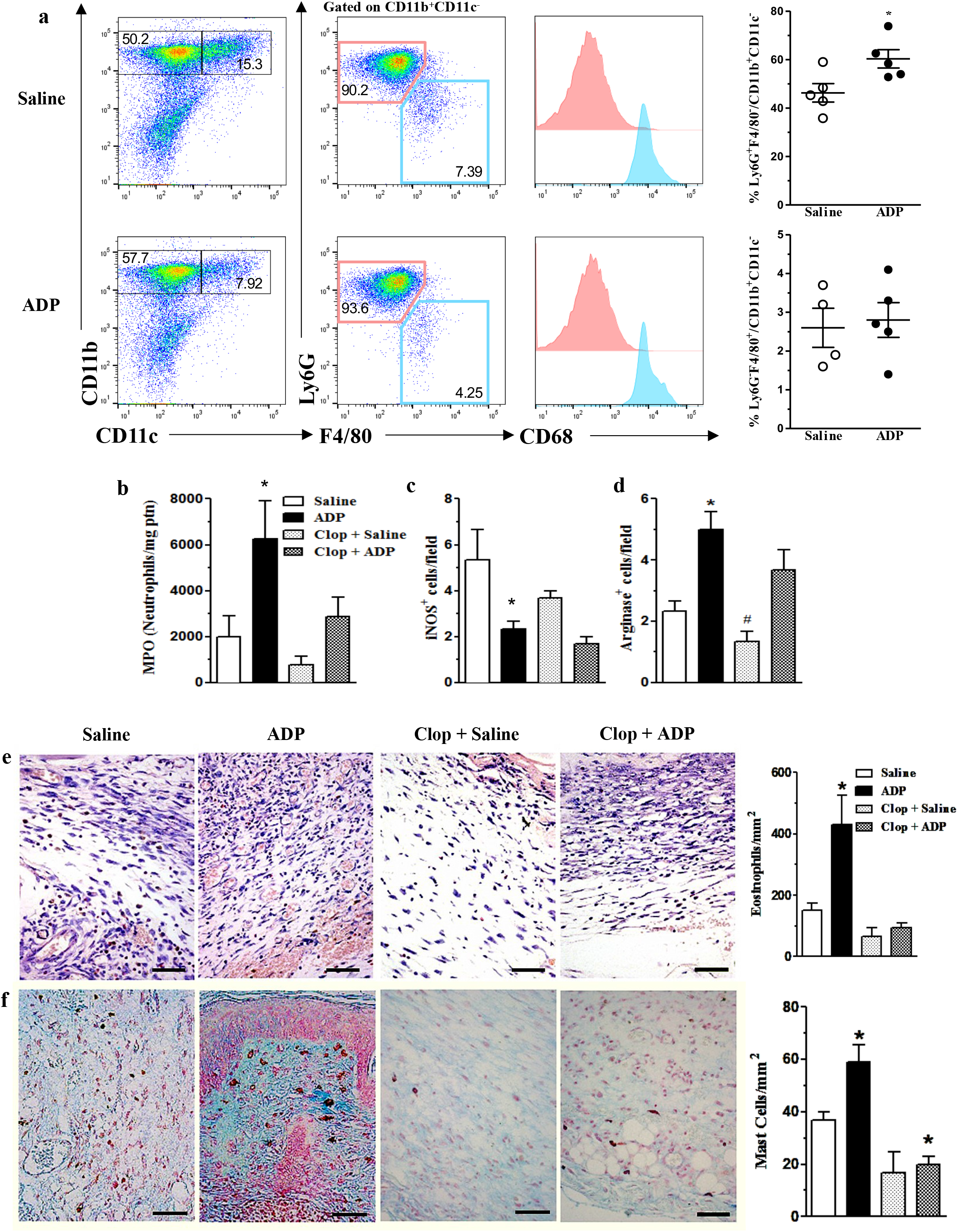
ADP-treated wounds present an increase of neutrophils, arginine^+^ cells, eosinophils and mast cells. Diabetic mice were submitted to excisional full-thickness wounding and, then, topically treated with ADP (30 µM/mouse) or saline every day for 7 days. Some mice were treated by gavage with Clop (5 mg/kg) 1 h before ADP or saline treatment, once a day for 7 days. (**a**) Wounds were harvested at day 7 after wounding and the cell suspensions were analyzed by flow cytometry. Dot plots (left) and graphs (right) show Ly6G^+^F4/80^-^ and Ly6G^-^ F4/80^+^ populations (gated on CD11b^+^CD11c^-^ cells) in the wounds. Graphs show the frequency of each cell population relative to gated live cells; for that, percentage of Ly6G^+^ [pink-neutrophils] or Ly6G^-^ [blue-macrophages] cells were multiplied by percentage of live CD11b^+^CD11c^-^ cells; intracellular CD68 staining confirm macrophage identity; data represent one experiment with n=4-5 mice per group; (**b**) Wounds were harvested at day 7 and prepared for myeloperoxidase quantification; n=6 per group and representative of three independent experiments; (**c**) Wounds were evaluated by immunohistochemistry for inducible nitric oxide synthase^+^ or (**d**) arginase^+^ cells at day 7 after wounding. *P<0.05 by Student’s t test, compared to saline-treated group and ^**#**^P<0.05 by Student’s t test compared to ADP-treated group; n=6 per group and data are representative of two independent experiments; (**e**) Skin histological sections and bar graph of wounds harvested at day 7 and stained with modified Sirius Red stain for eosinophil or (**f**) with Alcian Blue stain for mast cells. Scale bars=50µm. ^*^P<0.05 by Student’s t test, compared to saline-treated group, and ^**#**^P<0.05 by Student’s t test compared ADP-treated group. Data referred of one experiment with 5-6 mice per group.

Moreover, histological examination of the skin sections from ADP-treated wounds showed increased eosinophil and mast cell populations compared to saline-treated wounds. Clop administration did not modify eosinophil and mast cell numbers in the saline-treated wounds, although Clop impaired ADP-induced accumulation of both cells, indicating P2Y_12_ involvement (Figure 6e-f).

T cells are resident in normal human skin and participate in cutaneous immunosurveillance, contributing to skin homeostasis [29]. Thus, we evaluated T cell profile in the skin and wound-draining lymph nodes of diabetic mice after ADP treatment. Interestingly, the percentage of Treg cells (Foxp3^+^/CD4^+^/CD3^+^) was selectively reduced in the ADP-treated wounds, but not in the draining lymph nodes (Figure 7a).

**Figure 7.**
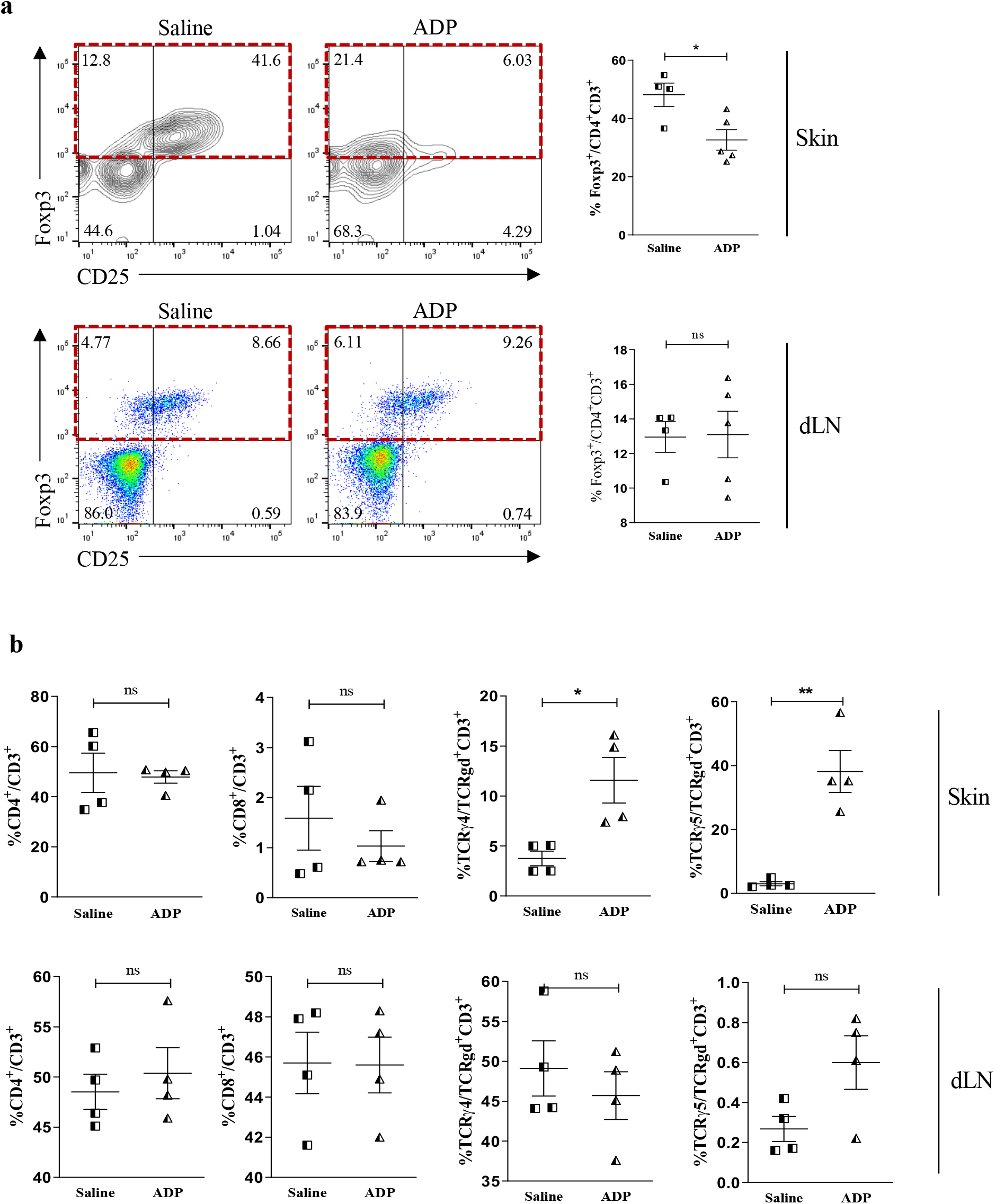
ADP treated wounds present a reduced population of Treg cells and an increase of Vγ4^+^γ*δ* T and Vγ5^+^γ*δ* T lymphocytes. Diabetic mice were submitted to excisional full-thickness wounding and then topically treated with ADP (30 µM/mouse) or saline every day for 7 days. Wounds and skin-draining lymph nodes were harvested at day 7 after wounding and the cell suspensions were analyzed by flow cytometry. Contour plots (top left) and dot plots (bottom left) and respective graphs (right) show the frequencies of (**a**) Foxp3^+^Treg cells (relative to CD4^+^CD3^+^population), (**b**) CD4^+^and CD8^+^ T cells (relative to total CD3^+^ lymphocytes), and (**c**) TCRγ4^+^ and TCRγ5^+^ cells (relative to total γδ^+^T lymphocytes). Data were expressed as mean standard error of the mean. *P<0.05 by Student’s t test compared to saline-treated group; n=4-5 per group, data are representative of two independent experiments, except for γd^+^T lymphocytes data that represent one experiment.

In parallel, ADP did not alter CD4^+^ and CD8^+^T cells frequencies in the skin and in the lymph nodes; however, ADP-treated wounds showed increased proportions of Vγ4^+^γ*δ* T cells and Vγ5^+^γ*δ* T cells. Again, no changes in T cell population were seen in the draining lymph nodes after wounding (Figure 7b).

### ADP accelerates wound closure only in diabetes condition

We also evaluated ADP effect on cutaneous ulcer induced by *Leishmania amazonensis* infection, and no improvement was observed (Figure 8a-b). These results indicate that ADP may be effective only in wounds due to metabolic diseases such as diabetes.

**Figure 8.**
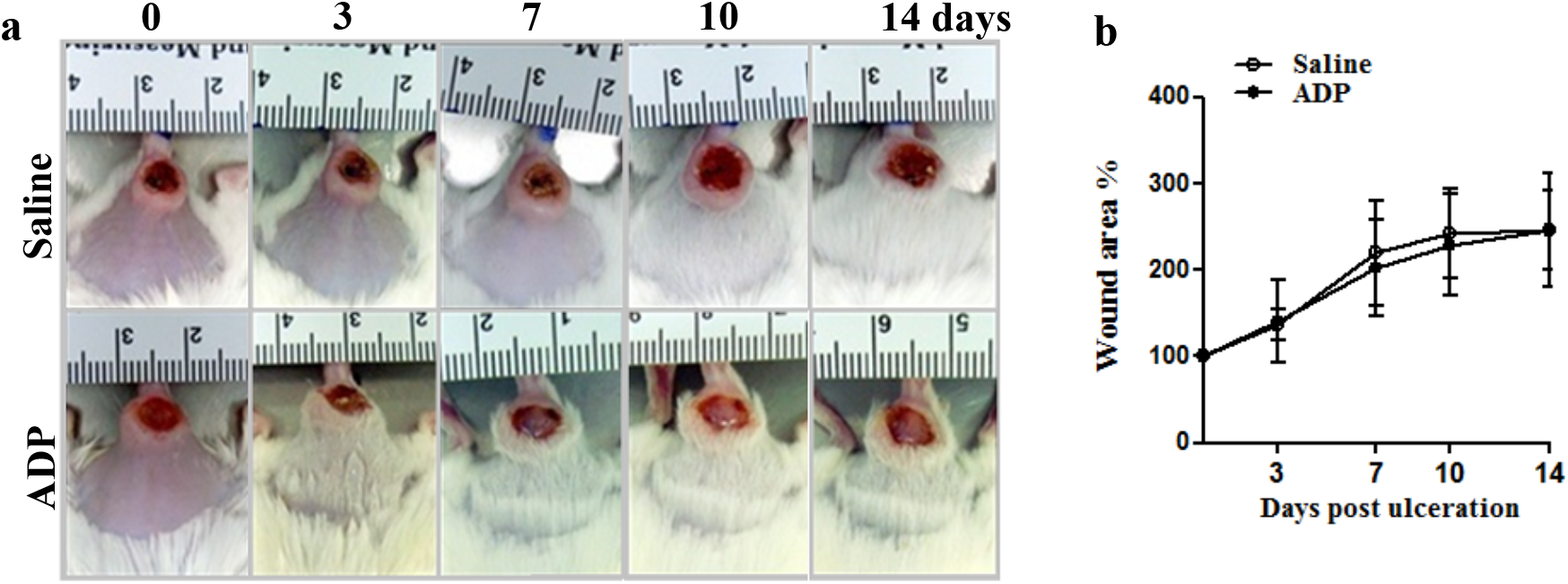
ADP did not accelerate wound healing of cutaneous leishmaniasis. BALB/c mice were intradermally inoculated with 10^6^ promastigote/mouse/50 uL (2×10^8^/mL). After wound ulceration, animals were topically treated every day with saline or ADP (0.38 µg/mouse) per 10 days. Open wound area was measured at days 0, 3, 7, 10, and 14. Representative images (**a**) and graph of cutaneous leishmania wound treated with saline or ADP (**b**). The areas at day 0 were considered 100 %, and the subsequent areas were proportional (%) to the initial wound area; n=7 per group.

## Discussion

Herein we demonstrate that ADP acts in accelerating skin wound healing in diabetic mice. Due to the large number of patients suffering from diabetes worldwide, presenting a poor life quality and high risk of complications as chronic wounds, we emphasize the importance to understand the pathophysiology of wound healing and to search for novel substances for wound treatment. ADP is an endogenous nucleotide involved in platelet aggregation, inflammation, and repair, without apparent side effects, being quickly metabolized. We have unequivocal evidence that ADP is a pivotal mediator for tissue repair and possibly a promising therapeutic agent for wound healing.

Controlled inflammation is essential for wound healing. The absence of inflammatory response or its exaggerated activation impairs wound healing towards the proliferative and remodeling/healing phases [30]. Our findings point ADP as a key modulator of cell activation, inflammation, and restoration of tissue integrity in diabetic mice wounds.

Adenosine diphosphate was able to improve wound healing only of diabetic mice. To note, ADP-treated wounds of diabetic mice heal at the same rate as wounds of non-diabetic healthy mice. Deficient wound healing in diabetic mice may be due to an insufficient ADP production, upregulation of ADP degradation, or ineffective expression/activation of purinergic receptors. Due to ADP liability its quantification was unfeasible. Thus, we moved to investigate the activity of enzymes that degrade extracellular nucleotides [31, 32]. Data from literature demonstrated that nucleotidases activity is increased in diabetic patients and in associated pathologies. Moreover, hydrolysis of adenine nucleotides is increased in platelets from diabetic patients [33,34]. Thus, we demonstrated a reduction in nucleotidase activity by ADP treatment in diabetic wounds. Furthermore, we also observed an increase on P2Y_1_ and P2Y_12_ receptors expression after ADP treatment. Taken together, both data may contribute to the positive effect of ADP in diabetic wounds.

Several strategies were used to certificate the ADP effect in wound healing of diabetic mice. Initially, we demonstrated that ATP, 5’AMP, adenosine, and pyrophosphate were not as effective as ADP. Additionally, an important role of endogenous ADP and P2Y_12_ in tissue repair was confirmed via Clop, since it impaired the beneficial effect of exogenous ADP on diabetic wounds, and also on non-diabetic wounds [26, 27, 34].

ADP’s receptors are expressed in important cells for tissue repairment [9, 26, 29], and, probably, this fact allowed ADP to improves tissue formation with less edema, collagen deposit, accelerated re-epithelization, and increased cell infiltration such as leukocytes and fibroblasts from the edge towards the center of the lesion. These cells characterize granulation tissue formation, crucial for healing process. The mentioned results place ADP as a pro-inflammatory and pro-resolution molecule, providing superior quality and organization in tissue formation.

Fibroblasts/myofibroblasts are contractile cells that approach the edges of the wounds and produce extracellular matrix, primarily collagen, the major mature scar component [35]. The increase of myofibroblasts population induced by ADP helps to explain the increase in collagen deposit and in dermal TGF-β^+^ cells. The shift from type III to type I collagen, triggered by ADP treatment, provides a more mature connective tissue and scar [35].

The balance of pro- and anti-inflammatory cytokines is essential for successful healing [36]. In our analyzes, the increment of IFN-γ levels, at day 3, as well as IL-10 and IL-13, at day 7 after wounding, suggests an anticipation in the shift of inflammation to resolution phase induced by ADP treatment. Also, the increased amount of TGF-β in the wound after ADP treatment corroborates an earlier resolution phase [37].

An intense inflammatory infiltrate in the wound was observed after ADP treatment. It is noteworthy that the inflammatory process, during normal wound healing, is characterized by spatial and temporal changes in leukocytes patterns. The well-defined chronology of these events is essential for ideal repair [38]. Tissue macrophages exposed to IL-4 and IL-13 are converted to a wound healing programmed cell [39]. The high concentration of IL-13 at day 7, together with a shift of macrophage phenotype from M1 to M2 after ADP treatment, corroborates to our hypothesis that ADP is a pro-resolution molecule.

Neutrophils can also manage the innate immune response during wound healing by regulating inflammatory response through the secretion of IL-10 [40]. Neutrophil accumulation and IL-10 secretion at day 7, induced by ADP, supports our hypothesis that ADP drives neutrophil activation towards a tissue repair profile.

Eosinophils infiltrate into wounds, often in proximity with fibroblasts, and stock and release TGF-β in rabbit cutaneous open wound models [41, 42]. Therefore, the eosinophils recruited by ADP may contribute to the accelerated wound healing via TGF-β production and collagen deposit, which correlates with improved tissue recovery. On the other hand, the role of mast cells in wound healing depends on the stimulus intensity [43]. In our diabetic model, the increased mast cell population, after ADP treatment, suggests a positive role for wound healing; but, other experiments are necessary to confirm this correlation.

Skin also hosts *α*β and γ*δ* T lymphocytes, which maintain skin homeostasis by modulating keratinocyte differentiation/re-epithelization, responding to infection, and regulating wound repair. A balance between regulatory T cell (Treg), Th17, and γ*δ* T cells orchestrates skin homeostasis [29]. The reduction of Treg cells and the increase of Vγ4^+^ and Vγ5^+^γ*δ* T cells in the wound, after ADP treatment, provide evidences of the recovery of epidermal barrier function and innate immunity response. Strikingly, mice deficient for γ*δ* T cells, including dendritic epidermal T cells (Vγ5^+^ TCR^+^), present a delay in wound healing and a defect to contain *S. aureus* infection [44, 45]. Vγ4^+^γ*δ* T cells migrate to dermis and epidermis after wounding and are a major source of IL-17A, which induces neutrophil migration, IL-1 and IL-23 production from epidermal cells to initiate local inflammation necessary for wound healing [46,47].

In conclusion, we demonstrated, for the first time, that ADP promotes skin homeostasis by inducing a brief and balanced inflammatory and immune response, followed by an adequate proliferative and remodeling phase, results summarized in the cartoon (Figure 9). Thus, exogenous ADP could be a new insight for therapeutic agents in diabetic wounds.

**Figure 9.**
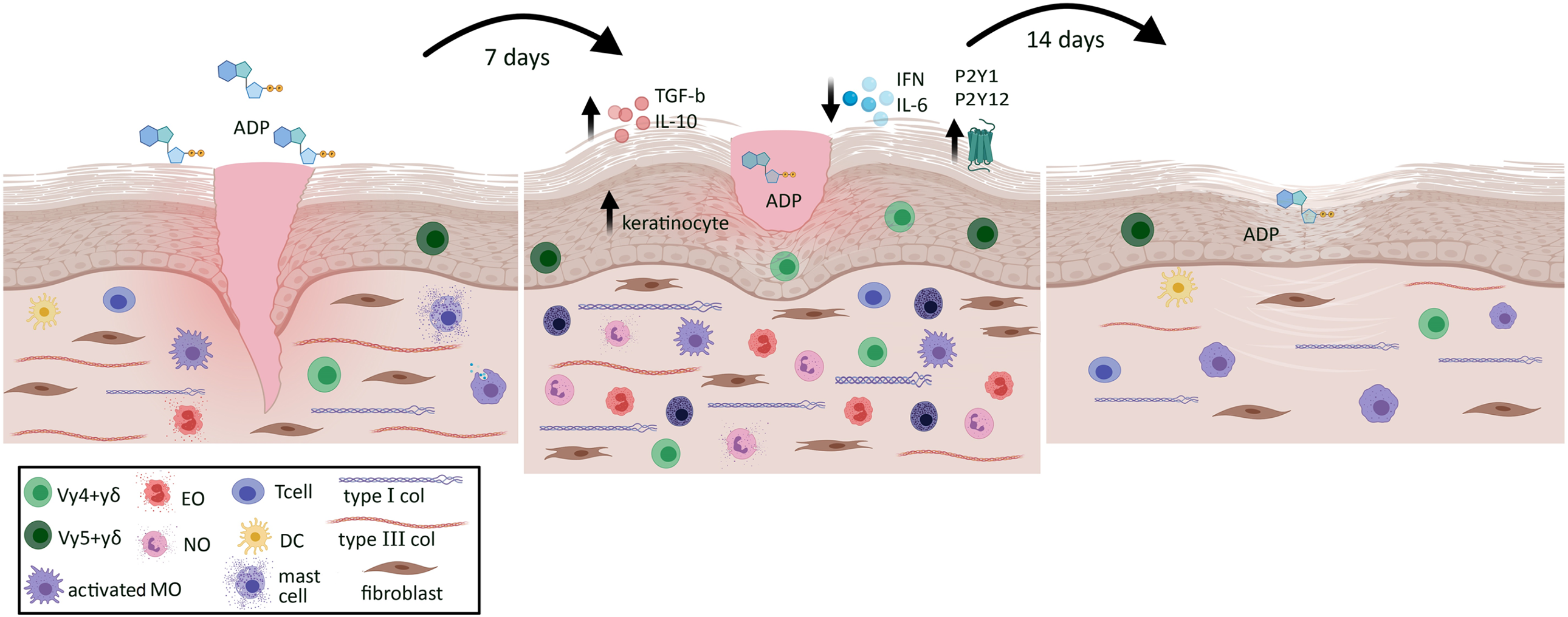
Summary of the pleiotropic effects of ADP on skin wound in diabetic mice. ADP topical instillation accelerates wound closure and improves tissue repair represented by type I collagen deposit and adequate reepithelization. The mechanisms seem to be via increase of neutrophils, eosinophils, mast cells, M2 macrophages, myofibroblasts and Vγ4^+^ and Vγ5^+^γ*δ* T cells in the wound, besides its ability in modulating cytokine release. ADP plays pivotal role within inflammation, proliferation and remodeling phases during skin tissue repair in diabetes wounds.

## Acknowledgement

We greatly acknowledge Ariane Rennó Brogliato for her contribution to the conception, design, analysis, and interpretation of the manuscript, and as a mentor to Paula Borges and Ingrid Waclawiak, it was of great significance. We thank Dr Isabela Ramos to kindly provide the pyrophosphate used in the *in vivo* wound healing experiment.

## Data availabitily

C.F.B. is the guarantor of this work and, as such, had full access to all the data in the study and takes responsibility for the integrity of the data and the accuracy of the data analysis. All data related to this paper may be requested from the authors.

## Authors’relationships and activities

The authors declare that there are no relationships or activities that might bias, or be perceived to bias, their work.

## Contribution statement

P.A.B., J.L.G. and I.W.: conceptualization, formal analysis, investigation, methodology, writing–original draft, and editing. J.F.B, V.S.F.J., F.S.L., and T.R.A: collected and analyzed the data for this work. E.M.S.: designed and performed the *Leishmania* experiment. CMT: performed and analyzed the histology data. R.C.S. and J.R.M.F: formal analysis and investigation of purinergic biology. C.P.: conceptualization formal analysis of γ*δ* T cells assay. J.S.N. and P.A.M.: conceived the project. F.C.: performed and analyzed all FACs data. C.M.: performed and analyzed IF data. C.C.: investigation, writing–review and editing. C.F.B.: conceived the project, formal analysis, investigation, project administration, resources, supervision, writing–original draft, writing–review and editing.

## Abbreviations

ADP: adenosine diphosphate
ATP: adenosine triphosphate
AMP: adenosine monophosphate
Clop: clopidogrel
α-SMA: α-smooth muscle actin
Treg: regulatory T cell.

## References

1. Gurtner GC, Werner S, Barrandon Y, Longaker MT (2008) Wound repair and regeneration. Nature 453:314–321. https://doi.org/10.1038/nature07039

2. Järbrink K, Ni G, Sönnergren H, Schmidtchen A, Pang C, Bajpai R, Car J (2016) Prevalence and incidence of chronic wounds and related complications: a protocol for a systematic review. Syst Rev 5(1):152. https://dx.doi.org/10.1186%2Fs13643-016-0329-y

3. Singer AJ, Clark RAF (1999) Cutaneous wound healing. N Engl J Med 341(10):738–746. https://doi.org/10.1056/nejm199909023411006

4. Stadelmann WK, Digenis AG, Tobin GR (1998) Physiology and healing dynamics of chronic cutaneous wounds. Am J Surg 176(2A Suppl):26S–38S. https://doi.org/10.1016/s0002-9610(98)00183-4

5. Tavares DMS, Dias FA, Araújo LR, Pereira GA (2009) Perfil de clientes submetidos a amputações relacionadas ao diabetes mellitus. Rev Bras Enferm 62(6):825–830.

6. Toscano MC, Sugita HT, Rosa MQM, Pedrosa HC, Rosa RS, Bahia LR (2018) Annual Direct Medical Costs of Diabetic Foot Disease in Brazil: A cost of illness study. Int J Environ Res Public Health 15(89):2–13. https://doi.org/10.3390/ijerph15010089

7. Van Battum P, Schaper N, Prompers L, Aprlqvist J et al (2001) Differences in minor amputation rate in diabetic foot disease throughout Europe are in part explained by differences in disease severity at presentation. Diabetic Medicine 28(2):199–205. https://doi.org/10.1111/j.1464-5491.2010.03192.x

8. Gachet C (2008) P2 receptors, platelet function and pharmacological implications. J Thromb Haemost 99(3): 466–472. https://doi.org/10.1160/th07-11-0673

9. Gendaszewska-Darmach E, Kucharska M (2011) Nucleotide receptors as targets in the pharmacological enhancement of dermal wound healing. Purinergic Signal 7(2):193–206. https://doi.org/10.1007/s11302-011-9233-z

10. Shen J, DiCorleto PE (2008) ADP stimulates human endothelial cell migration via P2Y1 nucleotide receptor-mediated mitogen-activated protein kinase pathways. Circ Res, 102:448–56. https://doi.org/10.1161/circresaha.107.165795

11. Battista AG, Ricatti MJ, Pafundo DE, Gautier MA, Faillace MP. ADP regulates lesion-induced in vivo cell proliferation and death in the zebrafish retina. J Neurochem 2009;111(2):600–613. https://doi.org/10.1111/j.1471-4159.2009.06352.x

12. Vadlamudi RVSV, Rodgers RL, McNeill JH (1982) The effect of chronic alloxana and streptozotocin-induced diabetes on isolated rat heart performance. Can J Physiol Pharmacol 60: 902–911. https://doi.org/10.1139/y82-127

13. im Walde SS, Dohle C, Schott-Ohly P, Gleichmann H (2002) Molecular target structures in alloxan-induced diabetes in mice. Life Sci 71:1681–94. https://doi.org/10.1016/s0024-3205(02)01918-5

14. Lanzetta PA, Alvarez LJ, Reinach OS, Candia OA (1979) An improved assay for nano-mole amounts of inorganic phosphate. Anal Biochem 100:95–97. https://doi.org/10.1016/0003-2697(79)90115-5

15. Lemos FS, Pereira JX, Carvalho VF et al (2019) Galectin-3 orchestrates the histology of mesentery and protects liver during lupus like syndrome induced by pristane. Sci Re 9(1):14620. https://doi.org/10.1038/s41598-019-50564-8

16. Kapoor M, Liu S, Huh K, Parapuram S, Kennedy L, Leask A (2008) Connective tissue growth factor promoter activity in normal and wounded skin. Fibrinogenesis Tissue Repair 1(1):3. https://doi.org/10.1186/1755-1536-1-3

17. Pavel M, Renna M, Park SJ et al (2018) Contact inhibition controls cell survival and proliferation via YAP/TAZ-autophagy axis. Nature Communications 9(1):2961. https://doi.org/10.1038/s41467-018-05388-x

18. Beyeler J, Schnyder I, Katsaros C & Chiquet M (2014) Accelerated Wound Closure In Vitro by Fibroblasts from a Subgroup of Cleft Lip/Palate Patients: Role of Transforming Growth Factor-a. PLoS One 9(10):111752. https://doi.org/10.1371/journal.pone.0111752

19. Brubaker AL, Schneider DF, Palmer JL, Faunce DE, Kovacs EJ (2011) An improved cell isolation method for flow cytometric and functional analyses of cutaneous wound leukocytes. J Immunol Methods 373:161–66. https://doi.org/10.1016/j.jim.2011.08.013

20. Meyerholz DK, Griffin MA, Castilow EM, Varga SM (2009) Comparison of histochemical methods for murine eosinophil detection in an RSV vaccine-enhanced inflammation model. Toxicol Pathol 37:249–55. https://doi.org/10.1177/0192623308329342

21. Carson LF, Hladik C (2009) Histotechnology: A Self-Instructional Text. American Society of Clinical Pathologists 188.

22. Devalaraja RM, Nanney LB, D. J, Qian Q, Yu Y, Devalaraja MN, Richmond A (2000) Delayed wound healing in CXCR2 knockout mice. J Invest Dermatol 115(2):234–44. https://doi.org/10.1046/j.1523-1747.2000.00034.x

23. Choi HS, Kim JW, Cha YN, Kim C (2006) A quantitative nitroblue tetrazolium assay for determining intracellular superoxide anion production in phagocytic cells. J Immunoassay Immunochem 27:31–44. https://doi.org/10.1080/15321810500403722

24. Yeh CJ, Chen CC, Leu YL, Lin MW, Chiu MM, Wang SH (2017) The effects of artocarpin on wound healing: in vitro and in vivo studies. Sci Rep 7(1):15599. https://dx.doi.org/10.1038%2Fs41598-017-15876-7

25. Burnstock G, Knight GE, Greig AVH (2012) Purinergic Signaling in Healthy and Diseased Skin J Investig Dermatol 132(3):526–546. https://doi.org/10.1038/jid.2011.344

26. Le Duc D, Schulz A, Lede V, Schulze A, Thor D, Brüser A, Schöneberg T (2017) P2Y Receptors in Immune Response and Inflammation. Adv Immunol 136:85–121. https://doi.org/10.1016/bs.ai.2017.05.006

27. Smith TM, Kirley TL (2006) The calcium activated nucleotidases: A diverse family of soluble and membrane associated nucleotide hydrolyzing enzymes. Purinergic Signal 2:327–333. https://dx.doi.org/10.1007%2Fs11302-005-5300-7

28. Xu F, Zhang C, Graves DT (2013) Abnormal cell responses and role of TNF-α in impaired diabetic wound healing. Bio Med Res Int 2013:754802. https://dx.doi.org/10.1155%2F2013%2F754802

29. Nestle FO, Di Meglio P, Qin J-Z, Nickoloff BJ (2009) Skin immune sentinels in health and disease. Nat Rev Immunol 9(10):679–691. https://dx.doi.org/10.1038%2Fnri2622

30. Cañedo-Dorantes L, Cañedo-Ayala M (2019) Skin Acute Wound Healing: A Comprehensive Review. Int J Inflam article ID 3706315. https://dx.doi.org/10.1155%2F2019%2F3706315

31. Vuerich M, Robson SC, Longhi MS (2019) Ectonucleotidases in intestinal and hepatic inflammation. Front Immunol 10:507. https://doi.org/10.3389/fimmu.2019.00507

32. Chia JSJ, McRae JL, Cowan PJ, Dwyer KM (2012) The CD39 adenosinergic axis in the pathogenesis of immune and nonimmune diabetes. J Biomed Biotechnol 2012:320495. https://doi.org/10.1155/2012/320495

33. Lunkes GI, Lunkes DS, Leal D et al (2008) Effect of high glucose levels in human platelet NTPDase and 5’-nucleotidase activities. Diabetes Res Clin Pract 81(3):351–357. https://doi.org/10.1016/j.diabres.2008.06.001

34. Cattaneo M (2006) P2Y12 receptor antagonists: a rapidly expanding group of antiplatelet agents. Eur Heart J 27(9) 1010–1012. https://doi.org/10.1093/eurheartj/ehi851

35. Werner S, Krieg T, Smola H (2007) Keratinocyte–Fibroblast Interactions in Wound Healing. J Investig Dermatol 127(5):998–1008. https://doi.org/10.1038/sj.jid.5700786

36. Kondo T, Ohshima T (1996) The dynamics of inflammatory cytokines in healing process of mouse skin wound: a preliminary study for possible wound age determination. J Leg Med 108(5):231–236. https://doi.org/10.1007/bf01369816

37. Ishida Y, Kondo T, Takayasu T, Iwakura Y, Mukaida N (2004) The essential involvement of crosstalk between IFN-gamma and TGF-beta in the skin wound-healing process. J Immunol 172(3):1848–1855. https://doi.org/10.4049/jimmunol.172.3.1848

38. Eming SA, Krieg T, Davidson JM (2007) Inflammation in wound repair: molecular and cellular mechanisms. J Investig Dermatol 127(3):514–525. https://doi.org/10.1038/sj.jid.5700701

39. Mosser DM, Edwards JP (2008) Exploring the full spectrum of macrophage activation. Nat Rev Immunol 8(12):958–969. https://doi.org/10.1038/nri2448

40. Zhang X, Majlessi L, Deriaud E, Leclerc C, Lo-man R (2009) Coactivation of Syk kinase and MyD88 adaptor protein pathways by bacteria promotes regulatory properties of neutrophils. Immunity 31(5):761–771. https://doi.org/10.1016/j.immuni.2009.09.016

41. Todd R, Donoff BR, Chiang T, Chou MY, Elovic A, Gallagher GT, Wong DT (1991) The eosinophil as a cellular source of transforming growth factor alpha in healing cutaneous wounds. Am J Pathol 138(6):1307–1313.

42. Basset EG, Baker JR, De Souza P (1977) A light microscopical study of healing incised dermal wounds in rats, with special reference to eosinophil leucocytes and to the collagenous fibers of the periwound areas. Br J Exp Pathol 58(6):581–605.

43. Tellechea A, Leal EC, Kafanas A et al (2016) Mast cells regulate wound healing in diabetes. Diabetes 65(7):2006–2019. https://doi.org/10.2337/db15-0340

44. Cho JS, Pietras E, Garcia N et al (2010) IL-17 is essential for host defense against cutaneous Staphylococcus aureus infection in mice. J Clin Investig 120(1 Supplement):17621773. https://dx.doi.org/10.1172%2FJCI40891

45. Girardi M, Lewis JM, Filler RB, Hayday AC, Tigelaar RE (2006) Environmentally responsive and reversible regulation of epidermal barrier function by gammadelta T cells. J Investig Dermatol 126(4):808–814. https://doi.org/10.1038/sj.jid.5700120

46. Li Y, Wu J, Luo G, He W (2018) Functions of Vγ4 T cells and dendritic epidermal T cells on skin wound healing. Front Immunol 9:1099. https://dx.doi.org/10.3389%2Ffimmu.2018.01099

47. Barros JF, Waclawiak I, Pecli C et al (2019) Role of Chemokine Receptor CCR4 and Regulatory T Cells in Wound Healing of Diabetic Mice. J Invest Dermatol 139(5):1161–70. https://doi.org/10.1016/j.jid.2018.10.039

