## Supplemental methods and data for "Adenosine diphosphate contributes to wound healing in diabetic mice through P2Y_1_ and P2Y_12_ receptors activation"

### Supplementary Materials

#### Supplemental Methods

##### Mice

Male Swiss and C57BL/6 male mice weighing 25-30 g were used for full-thickness excisional wound and ischemic wound models. The animals were obtained from Institute of Science and Technology in Biomodels (ICTB) in Oswaldo Cruz Foundation (FIOCRUZ). For cutaneous leishmaniasis lesion model we used male BALB/c mice weighing 25-30 g, obtained at Experimental Animal Facility from Microbiology and Parasitology Department in the Biomedical Institute in Federal Fluminense University. All animals were kept at least for one week in the same animal facility before use. All procedures described were reviewed and approved by the Ethics Committee for the Use of Animals of the Federal University of Rio de Janeiro (CEUA/UFRJ - processes: n° 093/15 and nº IMPPG 128/15) and the study was conducted in accordance with national guidelines for the care and use of laboratory animals.

##### Induction of diabetes mellitus

Diabetes was induced by i.v. injection of alloxan (65 mg/kg in saline) in mice fasted for 12 h[1,2]. Control mice (non-diabetic) were injected with saline. After administration of alloxan, animals were provided with food and water *ad libitum*. Diabetes was confirmed 7 days later when blood glucose concentration was at least 350 mg/dl, using the glucometer Accu-Check Active®. The glucose levels were still elevated (over 350 mg/dl) at day 30 after alloxan injection (data not shown).

##### Full-thickness excisional wound model

At day 7 after alloxan administration, animals were weighed and anesthetized with ketamine (112 mg/kg) and xylazine (7.5 mg/kg) by i.p. injections. Under anesthesia, a full-thickness excisional wound of 10 mm in diameter was created on the shaved dorsum of mice using biopsy punch. Wounds were treated once a day, every day, with topical application of different nucleotides (5'-adenosine monophosphate [AMP], ADP, 5'-adenosine triphosphate [ATP] - Sigma-Aldrich, St Louis, MO), adenosine (ADO - Sigma-

Aldrich), and pyrophosphate (Sigma-Aldrich) at 0.38 µg/ 30 µL/ mouse (30 µM - 15.4 µg/kg), or vehicle (30 µL of saline/mouse) for 5 or 14 days or until the day of sample harvest.

**Wound area quantification**

To determine the wound closure rate, the macroscopic wound area was evaluated by photos taken at days 0, 3, 7, 10, and 14 after wounding. Photos were taken at a standard distance using a tripod for the camera and they were analyzed using the ImageJ software. Wound closure data were expressed as a percentage of the initial wound area. In different set of experiments the skin sections were harvested at the same time points described above.

**Cutaneous leishmaniasis ulcer model**

To perform this assay, we used *Leishmania amazonensis* (MHOM/BR/75/Josefa) originally isolated from human cutaneous leishmaniasis, maintained at 26˚C in Schneider insect medium supplemented with 10% fetal calf serum (Gibco-BRL, Gaithersburg, MD) and 100 U/mL of penicillin and 100 µg/mL of streptomycin (Sigma-Aldrich). Promastigotes used at the stationary phase of growth (5-6 days of culture) were washed and resuspended in 0.01 M PBS pH 7.0 at $2 \times 10^8$/mL. Male BALB/c mice were intradermally inoculated with 50 µL ($10^7$ promastigotes/ mouse) in the shaved rump (at the tail base). After wound ulceration, animals were topically treated every day with saline or ADP (0.38µg/ mouse) per 10 days and the macroscopic wound area was evaluated.

**Treatments**

The administration of Clopidogrel® (Clop - 5 mg/kg) was performed daily by oral gavage, 1 h before ADP administration, for 14 days. Treatments with $P2Y_1$ (MRS 2179- 30 µM - 0.38 µg/ 30 µL/ mouse - Tocris, Bioscience, UK) and $P2Y_{12}$ (MRS 2395 - 30 µM, 30 µL/ mouse - Sigma-Aldrich) receptor antagonists, and with ATP diphosphohydrolase (apyrase - 6 U/mL, 30 µL/ mouse - Sigma-Aldrich) were performed topically for 14 days, 30 minutes before ADP administration.

**Histological procedures**

Wounds tissues were excised and fixed in 4% buffered formalin, dehydrated in an ethanol series, and paraffin-embedded sections (5-µm thick) were stained with hematoxylin–eosin (H&E) for general morphological aspect.

**Total collagen quantification**

Skin sample sections were stained with Picro-Sirius Red for total amount of collagen quantification. Seven micrometers thick sections of paraffin-embedded wound biopsy specimens were deparaffinized and rehydrated with distilled water, and then stained with a Picro-Sirius Red staining kit (Polysciences, Warrington, PA, USA), according to the manufacturer's instructions. The amount of collagen was determined by morphometric analysis using a quantitative imaging software (ImagePro Plus, version 4.5.1). Images were taken using a digital camera from a microscope at 40x magnification (Olympus BX53, Tokyo, Japan). Collagen fibers were stained in red, and the percentage of collagen per field was obtained by dividing the total area by fibrosis area.

**Ecto-nucleotidase activity measurements**

Ecto-nucleotidase activity was determined by the rate of inorganic phosphate (Pi) release. Wounds were removed and homogenized in 0.9% saline solution with 0.1% Triton X100 containing protease inhibitor cocktail (Roche, Mannheim, Germany). The suspensions were washed and incubated for 1 h at 25°C in 0.5 mL of reaction mixture containing, unless otherwise specified, 116 mM NaCl, 5.4 mM KCl, 5.5 mM d-glucose, 50 mM Hepes-Tris buffer, pH 7.2, and 5 mM ADP as substrate. The reaction was initiated by the addition of ADP and stopped by the addition of 1 mL of ice-cold 25% charcoal in 0.1 M HCl. This charcoal suspension was washed at least 20 times with 0.1 M HCl before use to avoid Pi contamination[19]. After the reaction, the tubes were centrifuged at 1500 g for 15 minutes at 4°C, and 0.5 mL of the supernatant was added to 0.5 mL of malachite green reactive mixture[20]. The absorbance of the released inorganic phosphate was measured by spectrophotometer at 660 nm. The ADPase activity was calculated by subtracting the nonspecific ADP hydrolysis measured in the absence of tissues samples. The concentration of Pi released in the reaction was determined using a standard curve of Pi for comparison and expressed as nucleotidase activity (nmol Pi x $h^{-1}$ x mg $ptn^{-1}$).

**Imunohistochemistry**

Seven micrometer thick sections of paraffin-embedded wound biopsy specimens were deparaffinized and hydrated and the slides were incubated with 10 mM sodium citrate. Endogenous peroxidase activity was blocked with 3% hydrogen peroxide. Slides were washed in TBS with 0.05% Tween 20 (Sigma-Aldrich), blocked with serum-free protein block (Dako, Santa Clara, CA), and immunostained with Vectastain® ABC HRP kit (Vector Laboratories, Burlingame, CA). Antibodies against type IA2 collagen (sc-28654 - 1:300, Santa Cruz Biotechnology, Santa Cruz, CA) and type III collagen (ab7778 - 1:100, Abcam), $P2Y_1$ (APR-009 - 1:200, Alomone Labs, Jerusalem BioPark, Israel), $P2Y_{12}$ (APR-012 - 1:200, Alomone Labs) transforming growth factor-β (TGF-β - ab66043 - 1:100, Abcam), arginase-1 (sc-166920 - 1:300, Santa Cruz Biotechnology), vascular endothelial growth factor (VEGF - sc-57496 - 1:300, Santa Cruz Biotechnology), Ki67 (ab15580-100 - 1:100, Abcam), and inducible nitric oxide synthase (iNOS - ab3523 - 1:100, Abcam) were used. Color was developed with 3,3-diaminobenzidine tetrahydrochloride (DAB - Vector Laboratories) and counterstained with hematoxylin. The data were expressed as number of positive cells per field. For collagen type markers, we developed a score method for the semi-quantification of collagen deposits performed by two different observers.

**Immunofluorescence**

For frozen sections, skin biopsy specimens were collected at day 7, embedded in optimal cutting temperature medium (OCT, Fisher Healthcare, USA), and 5 micrometer thick sections were cut on a cryostat (Leica, Germany). Then, slides were fixed with 4% paraformaldehyde in phosphate buffered saline and stained with antibody against $\alpha$-SMA (A-2547, 1:200, Sigma-Aldrich, USA). After incubation, cells were washed and incubated with Alexa Fluor-conjugated secondary antibodies (Invitrogen, Brazil), and nuclei were stained with DAPI (0.1 µg/ml in 0.9% NaCl). Specimens were examined with an Axiovert 100 microscope (Carl Zeiss, Germany) and images were acquired with an Olympus DP72 digital camera (Olympus, Japan). Control experiments with no primary antibodies showed only a faint background staining (data not shown).

**Cell proliferation**

Primary neonate dermal fibroblasts were plated (2 x $10^4$ cells) on coverslips in 24-well tissue culture dishes under standard culture conditions. After 24 h, 10 µL of BrdU (1 mg/mL - Thermofisher, Boston, MA, USA) was added to the culture medium. Cells were divided into 4 groups: ADP-treated cells (10, 30, or 100 μM) or untreated cells (DMEM-F12 medium 1% serum). After 24 h, the cells were fixed with 4% PFA for 30 minutes, washed three times with PBS 5 minutes each, and incubated with 2 M hydrochloric acid solution (HCl) for 30 minutes followed by incubation for 1 h with the solution containing 50 mM NaCl and 100 mM Tris hydrochloride, pH 7.5. Three washes were performed with PBS 10 minutes each, followed by three washes with 0.5% Triton X-100 in PBS for 10 minutes. Subsequently, a blocking solution containing 1% FBS in PBS was used for 30 minutes at 37ºC. Incubation with the anti-BrdU antibody (1:50) was performed for 1 h, followed by three washes with 0.5% Triton X-100 in PBS. Cells were incubated with anti-mouse secondary antibody at 1: 250 dilution (Alexa Fluor 555) for 1 h at 37ºC in the dark. Finally, cells were washed with PBS, and the coverslips were removed from the plate and positioned with the cells down into glass slides containing 5 μL of mounting medium for fluorescence with DAPI (Vector Laboratories). The images were captured using a fluorescence microscope and analyzed using ImageJ software. The proliferation was expressed as the percentage of $BrdU^{+}$ cells by total number of cells labeled with DAPI.

**Fibroblast migration**

Primary dermal fibroblasts were seeded onto six-well plates and grown until 90% confluence. After 24 h of serum starvation, cells were incubated with mitomycin-C (5 μg/mL - Sigma-Aldrich) for 2 h to avoid proliferation and then washed with PBS. Cell-free areas were created by disrupting the monolayers (scratch) with sterile pipette tip. Subsequently, medium was changed and cells were incubated with only medium or with 10, 30, or 100 μM of ADP until 24 h. Pictures of the scratched areas were taken immediately (0 h) after wounding and then at 6, 12, 18 and 24 h using an inverted microscope equipped with a digital camera (BEL Engineering - Monza, Italy). The scratched areas were measured using the ImageJ software, and the fibroblast migration was expressed as the % of area still open related to the initial area (time 0 h).

**Flow Cytometry**

Flow cytometry of wound tissues was performed as previously described[21]. Briefly, wounds tissue obtained from individual skin samples were digested by dispase enzyme solution (0.375 mg/mL - Roche Diagnostics) and enzyme cocktail (Sigma-Aldrich). Cells ($10^6$ cells/mL) obtained from each mouse were resuspended in PBS and blocked with Fc block (BD Biosciences Pharmingen, San Jose, CA). Different cell populations were identified using the antibodies in the table below.

| **Antibodies** | **Fluorochrome** | **Dilution** | **Clone** | **Source** |
|---|---|---|---|---|
| CD4 | BV605 | 1:200 | GK1.5 | Biolegend |
| CD4 | PE-Cy7 | 1:200 | GK1.5 | BD Biosciences |
| CD3 | BV421 | 1:200 | 17A2 | BioLegend |
| CD25 | APC | 1:100 | PC61.5 | eBioscience |
| FOXP3 | PE | 1:50 | FJK-16s | eBioscience |
| CD45.2 | FITC | 1:100 | 104 | BD Pharmingen |
| CD8 | PE | 1:200 | 53-6.7 | BD Pharmingen |
| TCRγδ | PerCP-cy5.5 | 1:80 | GL3 | Biolegend |
| TCRγ4 or (Vγ2) | APC | 1:80 | UC3-1046 | BD Pharmingen |
| TCRγ5 or (Vγ3) | FITC | 1:80 | 536 | BD Pharmingen |
| CD11b | FITC | 1:100 | Mac-1 | BD Pharmingen |
| CD11c | PE-Cy5 | 1:200 | N418 | eBioscience |
| F4/80 | PE-Cy7 | 1:200 | BM8 | BD Biosciences |
| CD68 | PE | 1:100 | FA-11 | Biolegend |
| Ly6G | Alexa 647 | 1:500 | 1A8 | Biolegend |
| Live/Dead | APC-Cy7 | 1:1000 | — | eBioscience |

Lymphocyte populations recovered from skin and draining inguinal, axillary, and brachial lymph nodes were analysed. For skin Treg cell analysis, samples were enriched by Percoll gradient for mononuclear cells. Samples were acquired with BD FACS Canto II (BD

Biosciences, San Jose, CA) and then analyzed with FlowJo software. The following gating strategy was used for T cell analysis: (a) FSC x SSC plot shows a gated region comprising leukocytes, (b) gated region on live cells, (c) gated region on $CD3^+$ cells, (d) dot-plot showing $CD4^+$, $CD8^+$, or $CD3^+TCR\gamma\delta^+$, and (e) gated TCRγ4 or TCRγ5 population among $CD3^+TCR\gamma\delta^+$. For Treg cell analysis: (a) FSC x SSC plot shows a gated region comprising leukocytes, (b) gated region on live cells, (c) gated region on $CD45^+CD4^+$ cells and (d) dot-plot of $CD25^+Foxp3^+$ cells. For granulocytes: (a) FSC x SSC plot shows a gated region comprising granulocytes, (b) gated region on live cells, (c) gated region on $CD3^+$ cells, (d) gated region on $CD11b^+CD11c^-$ and $CD11b^+CD11c^+$ cells, (e) gated regions on $Ly6G^+$ and $F4/80^+$ cells, and (f) histogram for CD68 expression on $Ly6G^+F4/80^-$ and $Ly6G^-F4/80^+$.

**Evaluation of eosinophilic infiltrate**

Sections were stained with modified Sirius Red for quantification of eosinophilic infiltrate[3]. Skin sections (5-µm thick) were deparaffinized and hydrated, washed in distilled water, stained with Harris hematoxylin, rinsed in 0.5% hydrochloric alcohol and washed in tap water. The sections were immersed in an alkaline (pH 8-9) Sirius Red solution and rinsed in water. Afterwards, sections were dehydrated in an ethanol series and cleared. Images were taken using a digital camera coupled to the microscope (Olympus BX53) at 40x magnification. Twenty fields were analyzed per wound/animal (n=3) and the data were expressed as number of eosinophils/$mm^2$.

**Evaluation of mast cells infiltrate**

Sections were stained with Alcian Blue for quantification of mast cell infiltrate. Tissue sections (5-µm thick) were deparaffinized and hydrated, rinsed in acetic acid (3%), stained with Alcian Blue (pH 2.5), rinsed again in acetic acid (3%), washed in water and stained with Neutral Red (0.5%), and washed briefly in distilled water. Afterwards, sections were dehydrated in an ethanol series and cleared. Images were taken using a digital camera from a microscope (Olympus BX53) at 40x magnification. Twenty fields were analyzed per wound/animal (n=3) and the data were expressed as number of mast cell/$mm^2$.

**Myeloperoxidase (MPO) activity assay**

To determine the MPO enzyme activity in the wounds, lesions were removed 7 days after wounding and homogenized in PBS containing protease inhibitors (Roche). The homogenates (50 mg of tissue/mL) were then centrifuged at 400 g for 10 minutes. The supernatant was resuspended in a solution of $Na_2PO4$ (80 mM) containing hexadecyltrimethylammonium bromide (HTAB - 0.5%) and centrifuged at 1,500 g for 15 minutes. For the MPO reaction, 60 µL of the supernatant was incubated with 4 µL 3,3,3,3 tetramethylbenzidine (TMB - 1.9 mg/ mL) and 40 µL $H_2O_2$ (1 mM) for up to 15 minutes. The reaction was stopped with sodium acetate (0.2 M) and the samples were evaluated by spectrophotometer at 630 nm. Neutrophil number was estimated by performing a standard neutrophil curve, using neutrophils obtained 6 h after peritoneal stimulation with 3% thioglycollate (more than 90% of neutrophils). Proteins were measured by Bradford method. The results were expressed as number of neutrophils per mg of protein.

**ELISA**

Seven days after wounding, lesions the wounds were removed (only the area not re-epithelized) and homogenized in PBS containing protease inhibitors. The homogenates were then centrifuged at 1,500 g for 10 minutes, and the supernatants were stored at -80°C freezer. The cytokine quantification (TNF-α, IL-10 and IL-13) was performed using PeproTech kits following the instructions of the manufacturer. Proteins were measured by Bradford method. The results were expressed as pg or ng of cytokine/mg of protein.

**Superoxide assay**

The superoxide production assay was performed by the nitroblue tetrazolium (NBT) reaction with reactive oxygen species resulting in formazan as final product[23]. Briefly, the wounds were removed at day 3 (only the area not re-epithelized) and homogenized in PBS containing protease inhibitors for the assay. The formazan formed was then solubilized in DMSO and potassium hydroxide (KOH - 2 M), and subsequently measured by ELISA reader (620 nm, Spectra Max-250, Molecular Devices). The results were expressed as µg of formazan/mg of protein.

**Cytometric Bead Array (CBA)**

The cytokines concentration in the wounds at days 3 and 7 was determined by flow cytometry using the kit Cytometric Bead Array (CBA) Mouse Inflammation (BD Biosciences, San Diego, CA) following manufacture's instructions. This CBA kit allows distinction of the following cytokines: IL-6, IL-10, MCP-1, IFN-γ, TNF-$\alpha$, and IL-12p70.

The sample processing and data analysis were acquired by FACSCalibur flow cytometer (BD Bioscences) and FCAP Array software, respectively. The results were expressed as pg or ng of cytokine/mg of protein.

**Western blotting**

Wound homogenates (30 mg of total protein) collected at day 7 were denatured in sample buffer (20 mM Tris-HCl + 1% SDS + 5% v/v β-mercaptoethanol + 10% v/v glycerol + 0.001% bromophenol blue) and heated in water (100°C) for 3 minutes. Samples were exposed to SDS-PAGE gel and the proteins were then transferred to PVDF membranes. The membrane was incubated in blocking solution TBS-T (0.05% Tween-20 in PBS) plus 5% BSA for 30 minutes, then incubated with primary antibody against α-SMA (1:1000 - Sigma-Aldrich) and β-actin (1:1000 - Cell Signaling, Danvers, MA) overnight at 4ºC. Secondary antibody (HRP conjugate) was incubated with the membrane at room temperature for 2 h. Following incubation, the membrane was washed three times with TBS-T for 5 minutes each. Immunoreactive bands were visualized using an enhanced chemiluminescence reagent (Amersham ECL, Biosciences) and pictures were recorded using Healthcare ImageQuant LAS 4000 (GE Healthcare Life Sciences, USA). The densitometry analysis was performed using ImageJ software and the results were expressed as the ratio of α-SMA/ β-actin (housekeeping).

**Statistical analysis**

Statistical differences in the wound closure experiments were determined using two-way ANOVA with the Bonferroni post-test in Graph Pad Prisms software for Windows. The significance of other experiments was determined by unpaired Student's t test.

#### Supplemental References

1. Vadlamudi, RVSV, Rodgers RL, McNeill JH (1982) The effect of chronic alloxan- and streptozotocin-induced diabetes on isolated rat heart performance. Can J Physiol Pharmacol 60(7):902-911. doi: 10.1139/y82-127

2. Im Walde SS, Dohle C, Schott-Ohly P, Gleichmann H (2002) Molecular target structures in alloxan-induced diabetes in mice. Life Sci 71(14):1681-1694. doi: 10.1016/s0024-3205(02)01918-5

3. Meyerholz, DK, Griffin MA, Castilow EM, Varga SM (2009) Comparison of histochemical methods for murine eosinophil detection in a RSV vaccine-enhanced inflammation model. Toxicol Pathol 37(2):249-255. doi: 10.1177/0192623308329342.

### Supplementary Figure

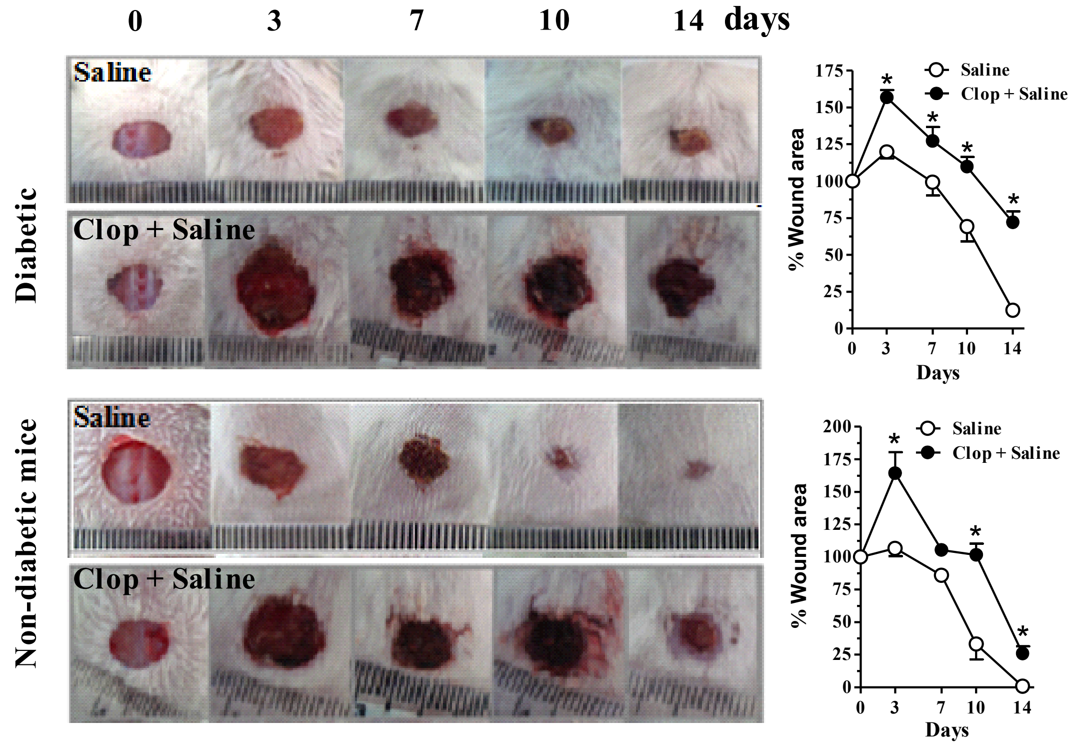


**Supplemental Figure 1.** Representative images and graphs of diabetic and non-diabetic mice submitted to excisional full-thickness wounding, and then treated by gavage with Clop (5 mg/kg) 1 h before topically saline (30 μL) treatment, both once a day for 14 days. Open wound area was measured at days 0, 3, 7, 10 and 14. The areas at day 0 was considered 100%, and the subsequent areas measured at different time-points were calculated as percentages (%) of the initial value. Data are expressed as mean ± standard error of the mean. *P<0.05 by Student's t test, compared to saline-treated diabetic or non-diabetic mice; n=7-10 per group.
